# MetaCell: analysis of single cell RNA-seq data using k-NN graph partitions

**DOI:** 10.1101/437665

**Authors:** Yael Baran, Arnau Sebe-Pedros, Yaniv Lubling, Amir Giladi, Elad Chomsky, Zohar Meir, Michael Hoichman, Aviezer Lifshitz, Amos Tanay

## Abstract

Single cell RNA-seq (scRNA-seq) has become the method of choice for analyzing mRNA distributions in heterogeneous cell populations. scRNA-seq only partially samples the cells in a tissue and the RNA in each cell, resulting in sparse data that challenge analysis. We develop a methodology that addresses scRNA-seq’s sparsity through partitioning the data into *metacells*: disjoint, homogenous and highly compact groups of cells, each exhibiting only sampling variance. Metacells constitute local building blocks for clustering and quantitative analysis of gene expression, while not enforcing any global structure on the data, thereby maintaining statistical control and minimizing biases. We illustrate the MetaCell framework by re-analyzing cell type and transcriptional gradients in peripheral blood and whole organism scRNA-seq maps. Our algorithms are implemented in the new MetaCell R/C++ software package.

## BACKGROUND

Single cell RNA-seq (scRNA-seq) is used extensively for discovery and identification of cell types, for characterizing transcriptional states within them, and for inference of continuous gene expression gradients linking these states. These phenomenological observations are used for creating cell type atlases, and as a starting point for analysis of different cellular processes, including differentiation, cell cycle and response to stimuli^1–9^ (reviewed in^10^). Key challenges in the analysis of scRNA-seq data are the discrete and variable nature of the cellular mRNA molecule census, and the sparsity of the scRNA-seq molecule count matrices aiming at its characterization. In mammals, only 10^5^-10^6^ copies of mRNA are present within each cell, representing regulated and stochastic transcriptional activities of over 20,000 genes^11^. At the same time, accurate quantitative models for the manner in which changes in the RNA abundance of specific genes distinguish cell types and cellular programs are lacking. In many cases RNA abundances can vary significantly due to biological noise, without a major effect on the cell’s current and future function. In other cases, genes with very low RNA molecule counts can have critical impact on the regulation of key programs. Finally, scRNA-seq technologies sample these stochastic mRNA pools sparsely, typically deriving between 10,000 unique molecule identifiers (UMI) from larger mammalian cells to less than 1000 UMIs for many important populations of smaller or quiescent cells^12^. Such sparse sampling inflicts additional variance on the estimates of RNA distributions in single cells. Effective scRNA-seq analysis must therefore take into account stochasticity occurring at multiple biological and technical levels^13,14^, and rely on large single cell cohorts to estimate functionally meaningful cellular programs while avoiding over-fitting and the introduction of model biases.

In the absence of sufficiently accurate parametric models for transcriptional variation in single cells, current scRNA-seq analysis methodologies rely heavily on non-parametric representations of the cells’ similarity graph. Combination of dimensionality reduction and embedding of single cells into 2D maps is used extensively for downstream visualization, clustering^15–18^, and inference of putative transition or differentiation gradients and the cells’ progression through them (termed *pseudotime*^19–26^). Many of the dimensionality reduction approaches are based on construction of K nearest neighbor (K-nn) graphs over single cells, and are therefore capable of modeling highly flexible non-linear trends in the data. By integrating multivariate information across many genes and cells, dimensionality reduction implicitly compensates for the sparsity and stochasticity of the data. In other cases such compensation is done explicitly, by either deriving clusters and pooling data from their cells^3,24,27^, or more recently by directly implementing strategies for data imputation^28–31^. When coping with sparsity involves global modeling of the underlying manifold, the analysis is at risk of becoming prone to modeling biases. Separating the two tasks is one way of avoiding this problem.

In this paper we introduce the notion of *metacells* and develop a methodology for inferring and using them. A metacell (abbreviated MC) is a group of scRNA profiles that are highly similar, ideally nearly indistinguishable from each other given some simplified parametric hypothesis on the sampled RNA distributions. In theory, a set of scRNA-seq profiles that are sampled from precisely replicated cellular RNA pools will be distributed multinomially with predictable variance and zero gene-gene covariance. Consequentially, any group of scRNA-seq profiles that are probabilistically consistent with sampling from the same RNA pool may be considered as an ideal metacell. Given a sufficient number of such homogenous profiles, we can estimate the distribution that generated them with high accuracy. In practice no two scRNA profiles sample exactly the same cell, since stochastic biological variation will diversify even the most functionally homogeneous cell populations. Still, we show below that in contemporary datasets sampling is sufficiently extensive to support partitioning of the cells into highly homogeneous metacells. In these cases, one can study the biological variation in the data using the rich set of inferred metacell states rather than the original sparse single cell profiles.

Metacells are building blocks for describing complex gene expression distributions with minimal parametric assumptions. Their sizes (typically in the order of 100 cells) are large enough to provide sufficient accuracy for transcriptional states estimation, and small enough to allow maximal flexibility for approximating quantitative gradients and dynamic biological processes across groups of metacells. By focusing on the characterization of disjoint local models without enforcing over them a global structure, the metacell approach accounts for data sparsity while minimizing modeling biases and smoothing artefacts. Finally, while the metacell model itself lacks any hierarchical or global component, it is possible to infer hierarchical structures by analyzing groups of metacells and their similarities.

We implemented tools for deriving metacells and analyzing scRNA-seq data using them in the new R/C++ package MetaCell. The utility of the approach was recently demonstrated for two different scenarios involving analysis of mammalian hematopoiesis differentiation^32^ and inference of cell type decompositions in comparative whole organism scRNA-seq^33,34^. Here we perform in-depth analysis of the model and its performance through re-analysis of datasets including 8,000 and 160,000 peripheral blood mononuclear cells (PBMC), and by dissecting two whole-organism single cell RNA-seq maps from two worm species. The data show that metacells approximate the expression distribution in a surprisingly accurate fashion, dissecting the dataset into truly homogenous local neighborhoods and providing quantitative building blocks for exploring the global expression distribution. We show how to diagnose outlier behaviors or transcriptional states that are not sufficiently sampled for effective approximation by metacells. We demonstrate how metacells can be used for robust analysis of transcriptional gradients, and how the approach avoids smoothing artefacts that are difficult to diagnose when analyzing the cell-cell similarity graph using common projection methodologies. Given the computational scalability of the method and the continuous increase in scRNA-seq dataset sizes, we believe metacells circumvent much of the difficulties associated with the sparsity of scRNA-seq data, and provide an attractive universal first layer of analysis on which quantitative and dynamic analysis can be developed further.

## RESULTS

### Overview of the MetaCell method

A MetaCell construction pipeline consists of the following stages (**Fig 1A**). First, feature genes are selected and used to compute a raw cell-to-cell similarity matrix *S*. Second, a balanced K-nn similarity graph *G* is constructed, connecting pairs of cells that represent reciprocally high-ranking neighbors. In contrast to a K-nn graph built directly from *S*, which can be highly non-symmetric, the graph *G* has more balanced ingoing and outgoing degrees. Third, *G* is subsampled multiple times, and each time the graph is partitioned into dense subgraphs using an efficient graph algorithm. The number of times each pair of cells co-occurred in the same subgraph is used to define the resampled graph *G*^*boot*^. After these three layers of cell-to-cell similarity matrix normalization, the metacell solution is derived using a graph partitioning algorithm applied to *G*^*boot*^.

**Figure 1:**
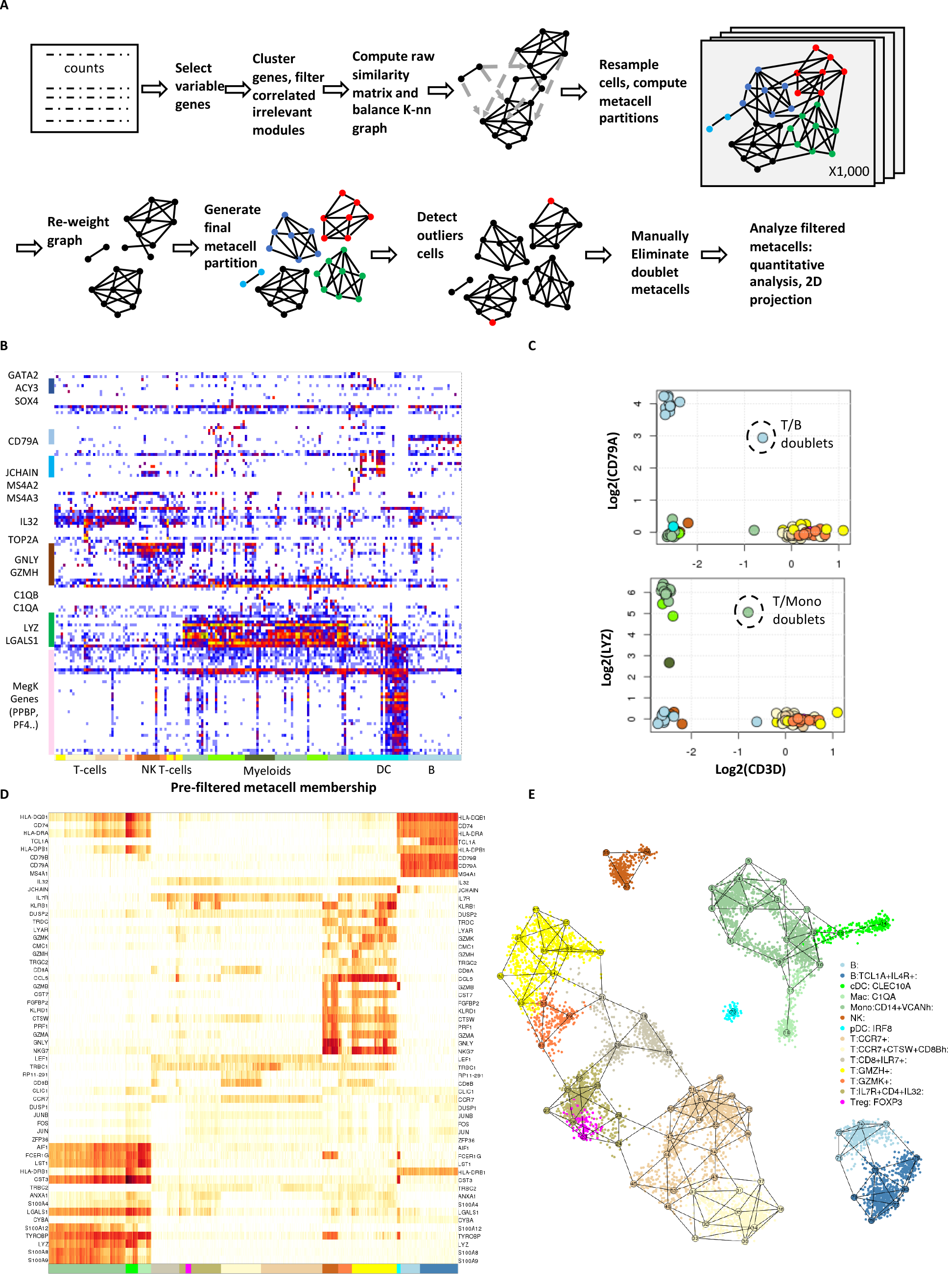
Metacell analysis of the PBMC 8K dataset. **A**) Schematics of the MC algorithmic pipeline. **B**) Outlier matrix showing color-coded number of UMIs per cells (columns) for which at least one gene (rows) was shown to be expressed significantly beyond its MC expected number of UMIs. Outlier cells are ordered according to the annotation of the MC containing them (bottom color-coded bars). **C**) Shown are log-fold-enrichment (lfp, methods) values for metacells, color-coded according to initial cell type annotation, comparing the T-cell marker (CD3D) to a B cell (CD79A) and myeloid (LYZ) markers. **D**) Heat map shows enrichment values for metacells (columns) and their maximally enriched gene markers. **E**) Shown is the MC adjacency graph (numbered nodes connected by edges), color-coded according to their cell type and transcriptional state annotation. Cells are shown as small color-coded points localized according to the coordinates of MCs adjacent to them. Fig S3 shows the adjacency matrix that was used to generate the projection.

After constructing metacells we perform validation, filtering and annotation as follows. First, outlier cells are detected and filtered using a simple test for gene overexpression compared to their metacell. Second, the metacells’ homogeneity is verified. This stage may result in splitting of a metacell into multiple metacells, but in practice this is seldom necessary. Third, metacells representing doublets are searched for and filtered in a supervised manner. Fourth, an expression profile and a list of marker genes, i.e. genes strongly enriched relatively to other metacells, are computed per metacell. Fifth, summary of adjacencies between metacells (based on adjacencies between their cells in the graph *G*) is used for clustering and high-resolution metacell annotation, such as cell type and subtype dissection. Finally, a 2D projection of the metacells and their constituting cells is generated for visualization purposes.

MetaCell is readily applicable as an R/C++ package and is scalable to large datasets. The full method and implementation details are given in the Methods section and the Appendix.

### Metacells eliminate outliers and reconstruct cell type structure in PBMC data

We illustrate the use of the MetaCell algorithm and pipeline (**Fig 1A**) through re-analysis of a small (n=8,276) dataset of PBMC scRNA-seq profiles sampled from a healthy donor and downloaded from the 10x website. In a pre-processing step we removed cells with less than 800 UMIs (**Fig S1A**) and several non-coding RNAs linked with stress or apoptotic signatures (“blacklisted genes”) (**Fig S1B**). We then applied the metacell construction pipeline as outlined above, using 816 high variance genes as features (**Fig S1C**, excluding ribosomal proteins) and deriving an initial set of 82 MCs following 1,000 resampling iterations using K=100. The MC outlier detection screen then identified 182 cells with at least one outlier gene (8 fold or more enrichment over the respective MC model) (**Fig 1B**, **Fig S2**). Most outlier cells showed potential doublet profiles, co-expressing genes associated with two different cell types. For example, this effect was notable in the association of a coherent megakaryocytic gene module (including PF4, PPBP and more genes) with signatures linked to other cell types. In fact, pure megakaryocyte expression profiles are very rare in the data and the MC outlier analysis highlights their identification (**Fig S2**). In addition to potential doublets, outlier cells also included representatives of rare cell types, including cells expressing progenitor markers (SOX4^35^) or eosinophilic markers (MS4A2, MS4A3^36^).

Doublet outlier cells are observed when two cell types are mixed rarely in the data, thereby contaminating a metacell associated with one cell type with a few mixed signatures. More frequent doublet scenarios can give rise to homogeneous doublet MCs, as we observed for two cases combining expression of T cell marker genes (e.g. CD3D) with either B cell (CD79A) or monocyte (LYZ) markers (**Fig 1C**). Following the removal of these two doublet MCs, we ended up with a model organizing 7,901 cells in 80 MCs (45-176 cells per MC, median size 95 cells) and marking 375 cells as outliers or doublets. This model was annotated using enriched gene markers (**Fig S3**), and visualized using a marker heat map (**Fig 1D**) and a 2D layout computed from the MC adjacency matrix (**Fig 1E**). This visualization organizes transcriptional states in the blood into clear cell type groups representing T, NK and B cells, monocytes/macrophages and DC populations. Within these cell types, the maps show additional structure. For example, T cells were organized into CD8+ effector states (marked by GZMH and additional genes), CD8+ pre-effector states (marked by GZMK+), CCR7+ CD8+ cells with variable degree of cathepsin-W (CTSW) expression, naïve CD8+ cells (IL7R+) and CD4+ cells showing some activation of Treg genes (FOXP3+). Overall, when sampling at a depth of 8,000 cells, the metacell analysis allowed for robust identification of cell types and initial modelling of gene expression distribution within them. Additional coverage can lead to refined modelling of transcriptional distributions within cell types as we shall demonstrate below, but first, we will use the basic model to evaluate some of the assumptions underlying the definition of metacells and their derivation.

### MetaCell graphs define a symmetrized and modular adjacency structure between MCs

MetaCell transforms the initial non-symmetric K-nn similarities into a balanced graph, and then analyses resampled partitions of this graph relations to yield a regularized the cell-to-cell similarity structure. This regularization gives rise to a variant of the original K-nn graph whose degrees are bounded by the sizes of cell type clusters in the data, and that shows low distortion between incoming and outgoing edges. The impact of this transformation on the degrees of the graphs generated for the PBMC dataset is depicted in **Fig 2A**, showing how initial balancing reduces the variance of in-degrees in the graph, and how the in- and out-degrees become more similar in the co-occurrence graph. Balancing is further shown to reduce spurious connectivity between cell types and increase the modularity of the MC graph. This can be demonstrated when analyzing MC adjacency matrices that summarize total connectivity between cells within each pair of MCs. Comparing raw K-nn, balanced, and resampled MC similarities (**Fig 2B** and compare **Fig S4**) shows for example connectivity from NK cells (MC #56) toward T cells and from pDCs (MC #70) toward multiple cell types in the raw matrix, which are eliminated in the balanced and resampled matrices. This comparison also highlights cases of myeloid MCs connecting a large group of monocyte MCs and cDCs (#15) or monocytes and macrophages (#17), that provide better separation with the more differentiated MCs in the balanced and resampled matrices. The resampled matrix in particular provides improved modularity within the large group of T-cell MCs, for example, grouping of CCR7+ T-cell MCs into distinctive clusters. In summary, in a typical scRNA-seq dataset the combination of abundant and rare states leads to an asymmetric K-nn structure linking rare cells with hubs within large clusters and resulting reduced graph. The MetaCell graph balancing procedure alleviates such effects, and we demonstrated here how this can lead to easier detection of cell type clusters and hierarchical structures.

**Figure 2:**
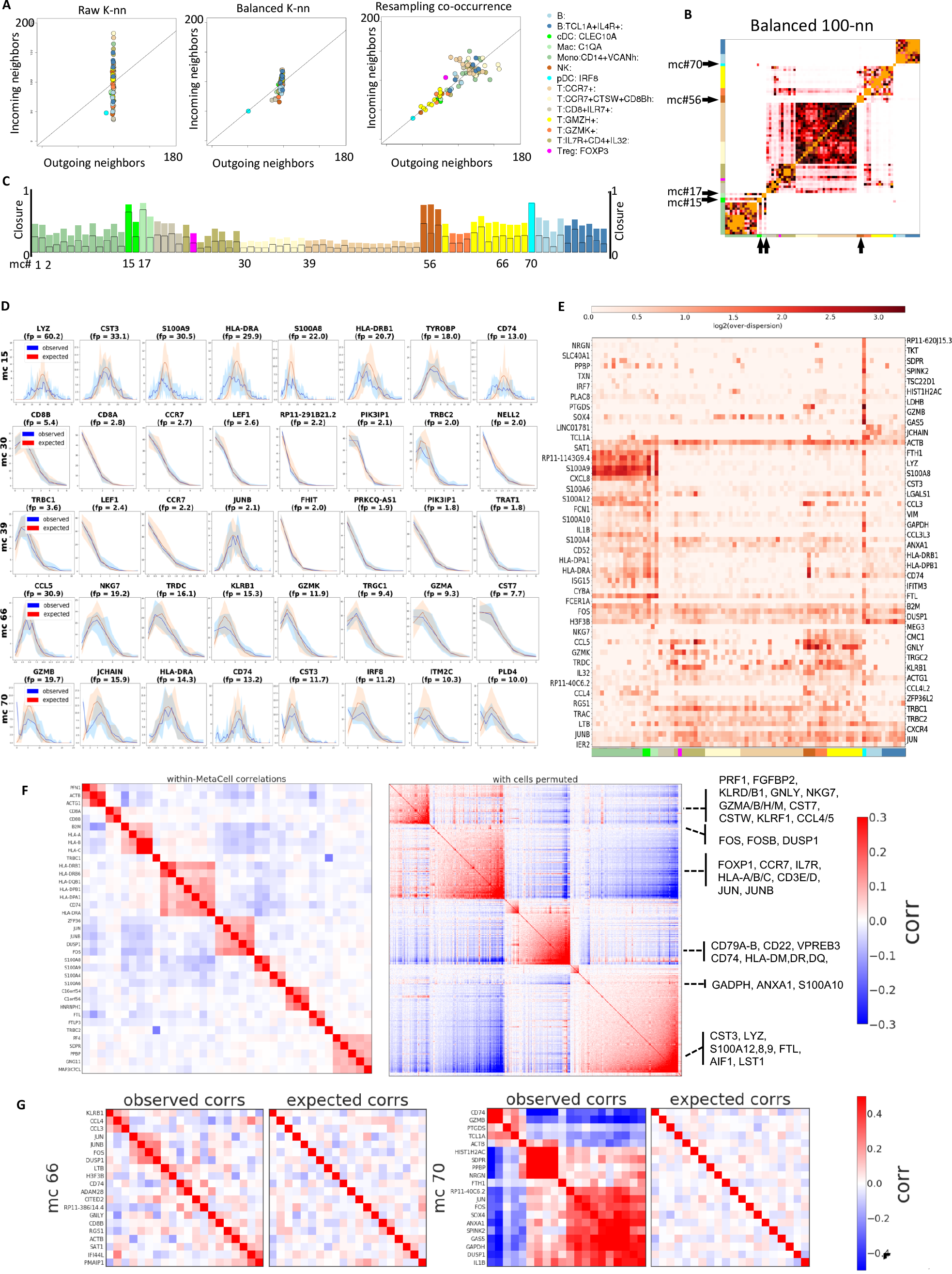
Evaluation of within-MC transcriptional homogeneity. **A**) Shown are the incoming and outgoing number of neighbors (or degree) per cell, averaged over metacells that are color-coded by cell type annotation as in Fig 1. The data represent the raw K-nn similarity graph (left), balanced MC graph (center), and resampled co-occurrence graph (right). **B**) Heat map summarizing the number of edges in the balanced MC graph that link two cells associated with different MCs. Similar matrices generated based on the raw and co-occurrence graphs are shown in Fig S4. **C**) Bar graph shows the closure per MC (fraction of intra-MC edges out of all edges linking cells in the MC). **D**) Observed (blue) vs predicted (red, based on a binomial model) distributions of downsampled UMI count per gene within MCs. For each of the 5 MCs depicted the plots show binomial fit for the top 8 enriched genes. Intervals give 10^th^ and 90^th^ percentiles over multiple down-samples of the cells within each metacell to uniform total counts. **E**) Over-dispersion of genes relative to a binomial model across genes and MCs. Colors encode ratio of observed to expected variance across genes (rows) and MCs (columns). Only genes and MCs manifesting high over-dispersion are shown. **F**) Residual within-MC correlation patterns compared with global correlation patterns. Within-MC correlation matrix (left) was computed by averaging gene-gene correlation matrices across MCs, where each matrix was computed using log-transformed UMIs over down-sampled single cell profiles associated with the respective MC. Global correlation matrix (right) was computed in the same manner, but following permutation of the MC assignment labels. For both matrices only genes manifesting strong correlations are shown. **G**) Examples of residual intra-MC correlated genes, showing observed correlations (Pearson on log-transformed down-sampled UMI counts) compared to correlations expected by sampling from a multinomial model. MC (#66) show weak residual correlations reflecting mostly stress genes. MC #70 show stronger residual correlations, reflecting residual intra-MC variation.

### Comparing metacells’ graph closure with their transcriptional homogeneity

To quantify the accuracy of the MC approximation to the similarity graph, we computed the fraction of K-nn similarities captured within each MC, which we refer to here as the MC’s *closure*. As shown in **Fig 2C**, the level of closure varies considerably between cell types. Distinct and low abundance cell types can show very high closure (up to 100%), while multiple MCs that cover abundant cell types show overall low closure (as low as 10% within-MC adjacencies, 20-30% within the three most linked MCs). Imperfect closure may suggest that the MC partition is suboptimal or, alternatively, that the K-nn local similarity structure in large and diffused cell types is covered by multiple, non-maximal but still homogeneous MCs. To test this, we compared the intra-MC UMI distribution to the distribution predicted by a simple multinomial model for specific genes and MCs (**Fig 2D**). We found that low closure MCs show high degree of consistency with the multinomial model, confirming their homogeneity. Interestingly, MCs with very high closure may show a reciprocal behavior, where additional high variance is present within K-nn consistent clusters (e.g. MC #70; note the bimodal distributions observed for most genes). This analysis highlights a key property of the MC partition: MCs are not maximal, and multiple highly similar MCs which are only weakly separated in the similarity graph can together approximate a larger cluster.

### Multinomial sampling explains most of the intra-MC UMI variance

Systematic screening for genes showing intra-MC over-dispersion (**Fig 2E**) provides a global view on the consistency of the PBMC MC cover with simple multinomial sampling. In this screening, MCs containing residual, non-homogeneous structure will be associated with many over-dispersed genes. For example, this analysis associates the dendritic cells MC #70 with over-dispersion of multiple megakaryocyte-associated and other genes. This suggests that these poorly sampled cell types show additional hidden structure and potential remaining outlier cells. The screening also reveals specific genes that are consistently over-dispersed across many MCs, such as the early-immediate response gene module (including the transcription factors JUN, JUNB, FOS). This over-dispersion is consistent with variable levels of activity of this pathway in multiple cell types, perhaps representing technical experimental stress. Other genes are over-dispersed in a cell-type specific fashion, for example cytotoxic (GNLY, CCL5) genes in NK and T subtypes, and MHC-II and LYZ in myeloid cell types. These highly-expressed genes may be incompatible with a simple multinomial sampling model, and their analysis may necessitate assuming prior biological variance to allow for over-dispersion. Beyond these specific examples, however, intra-MC distributions for the entire gene set (including genes that were not used as features for defining similarities) are generally well approximated by Poisson sampling with no zero-inflation (**Fig S5**). Together the data shows that the degree of residual, intra-MC over-dispersion is relatively low in the PBMC MC cover, so that the variance of most genes is accounted for by a model assuming partition of cells into MCs from which UMI’s are multinomially sampled.

Analysis of intra- and inter-MC gene-gene covariance (**Fig 2G**) provided an additional avenue for diagnosing structure within and between MCs. We observed persistent intra-MC correlations between a limited set of genes, including the over-dispersed modules of early-immediate genes, MHC class II genes, S100 genes as well as a correlated gene set including actin-related genes (ACTB, ACTG1, COTL1, PFN1). We did not observe strong intra-MC correlations of cytotoxic and many other functional genes. The scarcity of strong intra-MC gene-gene correlations (see for example **Fig 2G**-MC #66) suggests that little residual structure remains within the MCs, and that the dataset is well summarized by the MC profiles. In the few cases where intra-MC correlations are observed (**Fig 2G**-MC #70), they indicate the need for a more flexible intra-MC modelling, or alternatively call for deepening the dataset with more cells defining the transcriptional states underlying the MC.

### Metacells are accurate local approximations of the expression manifold

All approaches for analysis of scRNA attempt to describe aspects of the expression manifold, each relying on different assumptions. MetaCell generates a high-resolution partition of the data, thereby focusing on approximating it locally. We tested the quality of this approximation using a cross-validation scheme, in which we predict the expression of each gene using a MetaCell model trained on data from which the gene was left out. **Fig 3A** illustrates the outcome of such prediction, showing accurate prediction for highly expressed genes and lower accuracy for low-UMI counts, for which sampling variance is high. We wanted to compare these predictions to those obtained using the models that underlie commonly used approaches for scRNA-seq analysis. To this end, we computed the cell-to-cell similarity matrices inferred by Seurat’s^15^ PCA-based approach and by a diffusion strategy as implemented in MAGIC^28^. We also included in the comparison the similarity matrix *S* initiating the MetaCell balancing process. For all similarities we employed the same cross-validation scheme that was applied to the MetaCell model, and computed local predictions by averaging 50 nearest neighbors for Seurat and *S*, and weighting all cells by their similarities for MAGIC (See Methods for a complete description).

**Figure 3:**
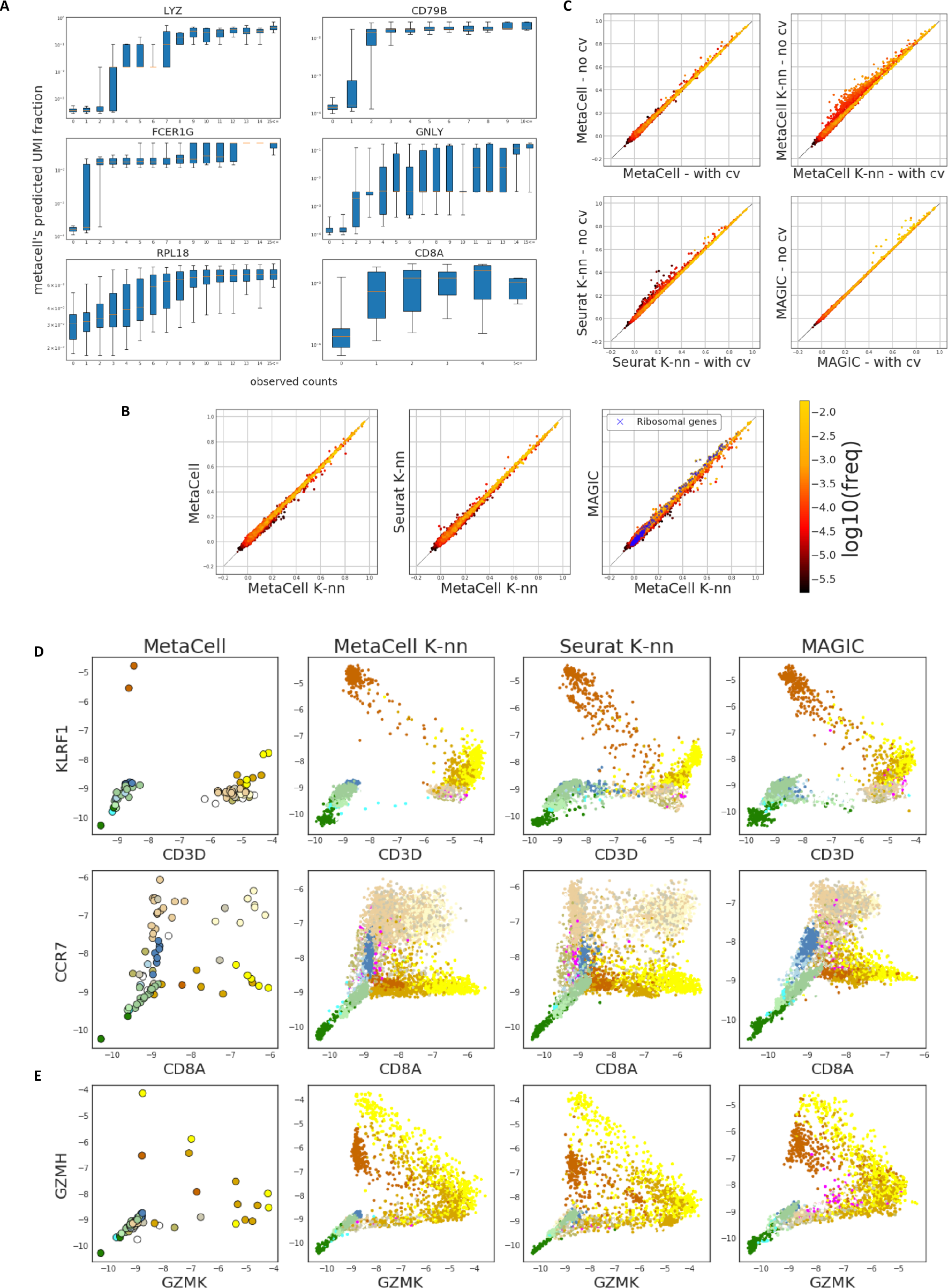
MCs robustly approximate the expression manifold. **A**) Boxplots show the distribution of predicted (using MC pool frequencies) UMI fraction per cell stratified according to observed number of UMIs in down-sampled single cells. **B**) Shown are per-gene Pearson correlations between predicted and observed gene frequencies for genes, color coded according to the gene’s frequency across all cells. In all cases predictions are generated using a 100-fold cross validation scheme (see Methods for exact description of the procedure and the strategies compared). Predictions using K-nns over raw MC similarities (a different neighborhood per cell consisting of its k most similar neighbors) is used as reference (X axis). It is compared to strategies defining cell neighborhoods using MCs (fixed disjoint grouping of cells), K-nn over Seurat distances, and MAGIC distances (weighted neighborhood according to diffusion distances). **C**) Similar to panels in (B) but comparing accuracy with and without applying cross validation. Points above the diagonal represent potential over-fitting. **D-E**) Per-MC (left most column) or smoothed per-cell (all other columns) expression values for pairs of genes, portraying putative transcriptional gradients.

Differences in prediction accuracy should reflect the different similarity measures employed by each method as well as the effect of disjoint partitioning applied in MetaCell. In theory, the partitioning strategy should provide less modelling flexibility compared to approaches that compute cell-specific neighborhoods. The latter effect should be particularly noticeable when several MCs discretize a continuum, such as differentiation trajectory. In practice, we observed relatively mild differences between the different approximations (**Fig 3B**), with very few genes losing accuracy when MCs are used. Moreover, analysis of the gain in accuracy when including all genes in the models (**Fig 3C**) suggested that MetaCell is significantly less exposed to over-fitting than the K-nn approaches. The diffusion-based smoothing approach showed minimal overfitting, but also loss of accuracy (**Fig 3C**). Overall, the nearly multinomial intra-MC UMI distribution observed above and the minimal loss of predictive power entailed by the MetaCell disjoint partition, together suggest that MCs succeed in capturing biological variation while reducing the residual intra-MC variation to sampling noise.

### Metacells avoid artefactual gradient effects

We showed that the cell partitioning induced by MetaCell does not decrease local approximation accuracy and that, in fact, it even reduces the model’s tendency to over-fit the data. We speculated that another advantage of partitioning would be robustness to over-smoothing. The discussion about over-smoothing recently arose in the context of evaluating scRNA-seq imputation methods, i. e. methods that use the covariance patterns measured across multiple cells and genes to refine per-gene, per-cell measurements (reviewed here^37^). Most imputation methods are local in the sense that they impute gene expression for a cell using its inferred neighborhood. It has been observed^29,38^ that in some cases imputation tends to enforce spurious proximities between cells, which in turn manifest as artefactual gradients, i.e. discrete states pertaining to be a series of cells gradually modulating expression of certain genes along a temporal process or a spatial axis. While over-smoothing is detected directly when evaluating imputation methods, it is in fact a potential concern with any model regardless of its downstream application, and stems from the manner in which cell-cell similarities are defined.

We evaluated the susceptibility of the MetaCell model to over-smoothing using the expression predictions obtained in the previous section (the version without cross-validation), comparing the different similarity structures included in that experiment. Our results support the robustness of MetaCell to artefactual gradients (**Fig 3D**). For example, NK cells are known to be characterized by high levels of KLRF1, but do not express the T-cell classical marker CD3 (**Fig 3D**, top). Smoothing based on K-nn similarity structures (MetaCell’s K-nn or Seurat’s) or on diffusion similarities (MAGIC’s) gives rise to phantom gradients that can be interpreted erroneously, for example, as supporting differentiation of NK to T-cells or vice versa. The MC statistics generate a much less detailed, but likely more realistic map of joint CD3D/KLRF1 expression. Similar phantom gradients are observed when analyzing CCR7+ CD8+ and CCR7+ CD8-cells (**Fig 3D**, bottom). On the other hand, the MC model does reveal expression gradients in cases where sampling adequately supports them, such as in the trade-off expression of GZMK+ and GZMH+ in T-cells (**Fig 3E**). These quantitative gradients are refined in the denser data set we analyze below.

### Dissecting complex cell type hierarchies with Metacell

We tested the scaling of MetaCell to datasets consisting of a large number of cell types and high variability in the total number of UMIs per single cell. To this end, we revisited two whole-organism scRNA-seq studies dissecting C. elegans (*Caenorhabditis elegans*)^39^ and Planaria (*Schmidtea mediterranea*)^40^. For C. elegans, we compared the derived MC partition (349 MCs) (**Fig 4A**, **Fig S6**) to the published model grouping cells into 27 major cell types (**Fig 4B**). We observed a high degree of consistency between the two models in classifying the major cell types, with higher resolution in dissecting cell types into subtypes using MCs (e.g. for body wall muscles, seam cells and more). Importantly, we observed a large number of cells labeled originally as “unclassified” or “unclassified neurons/glia” that were organized within coherent MCs. Some of these MCs were dominated completely or almost completely by unclassified cells. Moreover, we observed a negative correlation between the median number of UMIs per cell in a metacell and the fraction of unclassified cells within it (**Fig 4C**). Comparing the number of UMIs per cell within MCs showed consistently lower UMI counts for unclassified cells (**Fig 4D**). The transcriptional specificity of MCs containing large fractions of unclassified cells was uniformly high, as confirmed by observation of co-expression of specific transcription factors and genes within such MCs (**Fig 4E**). Similarly, Metacell analysis of the rich whole-organism cell type map of Planaria showed extensive consistency between the MC partition (564 MCs) and the iterative and highly supervised clustering analysis (512 clusters) used to annotate the original map (**Fig S7**). In summary, while MetaCell is not designed to perform clustering in its classical sense, a metacell partition facilitates robust and sensitive cell type mapping of scRNA-seq data, in particular when gene expression and cell type sizes are extremely heterogeneous.

**Figure 4:**
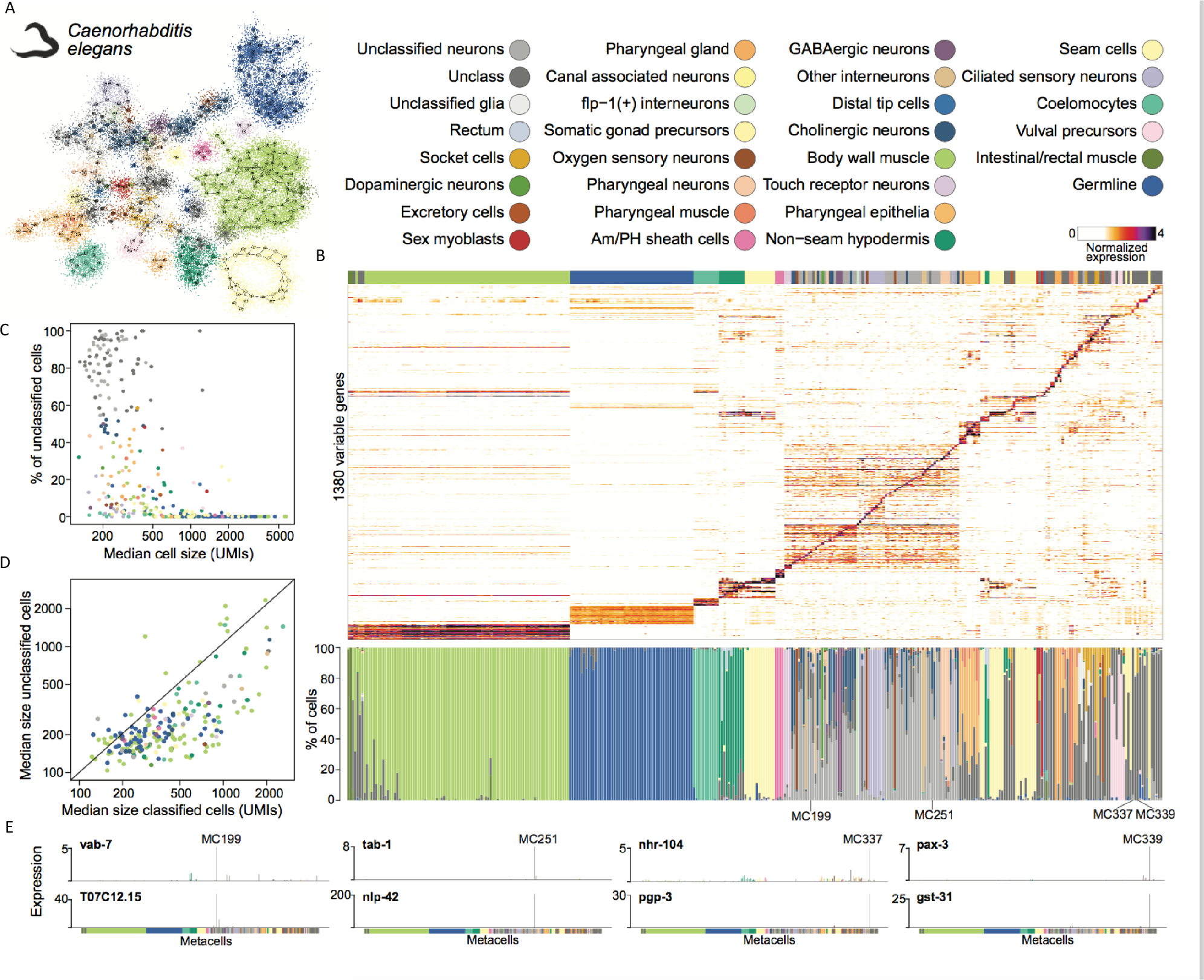
MC analysis of a whole-organism single-cell dataset. **A**) 2D projection of *C. elegans* metacells and single cells, color-coded according to the most frequent cell type based on the classification from Cao et al. **B**) Top - normalized expression of 1,380 highly-variable genes across 38,159 *C. elegans* single cells (columns), sorted by metacell. Bottom - barplot showing for each metacell the single-cell composition of the different originally classified cell types. **C**) Relationship between the metacell median cell size (UMIs/cell) and the fraction of cells originally labeled as “unclassified” in Cao et al. **D**) Comparison of the median sizes (UMIs/cell) of originally unclassified cells versus classified cells in each metacell. **E**) Expression (molecules/10,000 UMIs) of selected marker transcription factors (top row) and effector genes (bottom row) across all metacells, supporting high transcriptional specificity for four examples of metacells containing a high fraction (>80%) of originally unclassified cells.

### High-resolution analysis of inter- and intra-cell type states in the blood

We next tested the scaling of the MetaCell algorithmic pipeline when applied to datasets sampling deeply a relatively small number of cell types by analyzing RNA from 160K single blood cells, including 68K unsorted PMBCs and 94K cells from ten different bead-enriched populations^41^. We hypothesized that, with increased number of cells, we could derive MCs with enhanced quantitative resolution and increased homogeneity, thereby allowing a more accurate identification of regulatory states and differentiation gradients in the blood. We derived a model organizing 157,701 cells in 1,906 metacells, identifying 4,475 cells as outliers. **Fig 5A** summarizes the similarity structure over the inferred MCs, indicating partitioning of the dataset into T cells, NK cells, B cells, myeloid cells, megakaryocytes and progenitor cells. In-depth analysis of the emerging cluster and sub-cluster structure in this matrix allowed us to identify groups of related MCs for further analysis, in many cases providing us with the ability to zoom into transcriptional programs (cell groups numbered 1-13 on **Fig 5A**) within large scale clusters that were identified in the global metacell 2D projection graph (**Figure 5B**). Visualization of genes that were specifically enriched in such programs demonstrate both bimodal markers and putative quantitative gradients organizing MCs within and between types (**Fig S8**). For example, we observed the correlated (and bifurcated) intensity of CD8A and CD8B expression in cytotoxic and memory T-cells, the variable MHC-I expression (HLA-A,HLA-C) in different cell sub-types (group (6)), variable levels of granzyme K and granzyme H expression along a putative cytotoxic gradient of CD8+ cells (groups (1),(3)), and a group of MCs expressing cathepsin W and CCR7+ but without the cytotoxic gene module (group (5)). The analysis of specific gene families (see **Fig S9**) illustrates how multiple effector genes are activated in different cell types in a convergent fashion (**Fig S9A**). Analysis of transcription factor expression across the different subtypes (**Fig S9B**) provided an initial blueprint for the regulatory mechanisms defining the observed transcriptional states. Importantly, the integration of different sorting batches allowed for enhanced resolution in several hematopoietic lineages, in particular CD34+ progenitor cells (Fig 5A, group (11)). Nevertheless, all MCs within the non-progenitor cell types represented a balanced mixture of sorted and non-sorted batches (**Fig 5C**). We note that the metacells produced by MetaCell’s specialized partition algorithm cannot be reproduced by conventional clustering, at least when used naively. We demonstrate this by clustering the PBMCs with Seurat using parameters that force fine clustering, generating 817 clusters. As shown in **Fig S10A**, the MC partition is consistent with these fine clusters at the level of the coarse-grained cell types, but not at higher resolutions. The fine clustering solution generates clusters that are likely to be overfitting specific genes (**Fig S10B**). In summary, for the densely covered, multi-batch 160,000 PBMC datasets, MetaCell provides analysts with a platform for distinguishing cell types and their internal hierarchies, and a robust scheme for characterizing quantitative expression gradients with guarantees against spurious smoothing effects.

**Figure 5:**
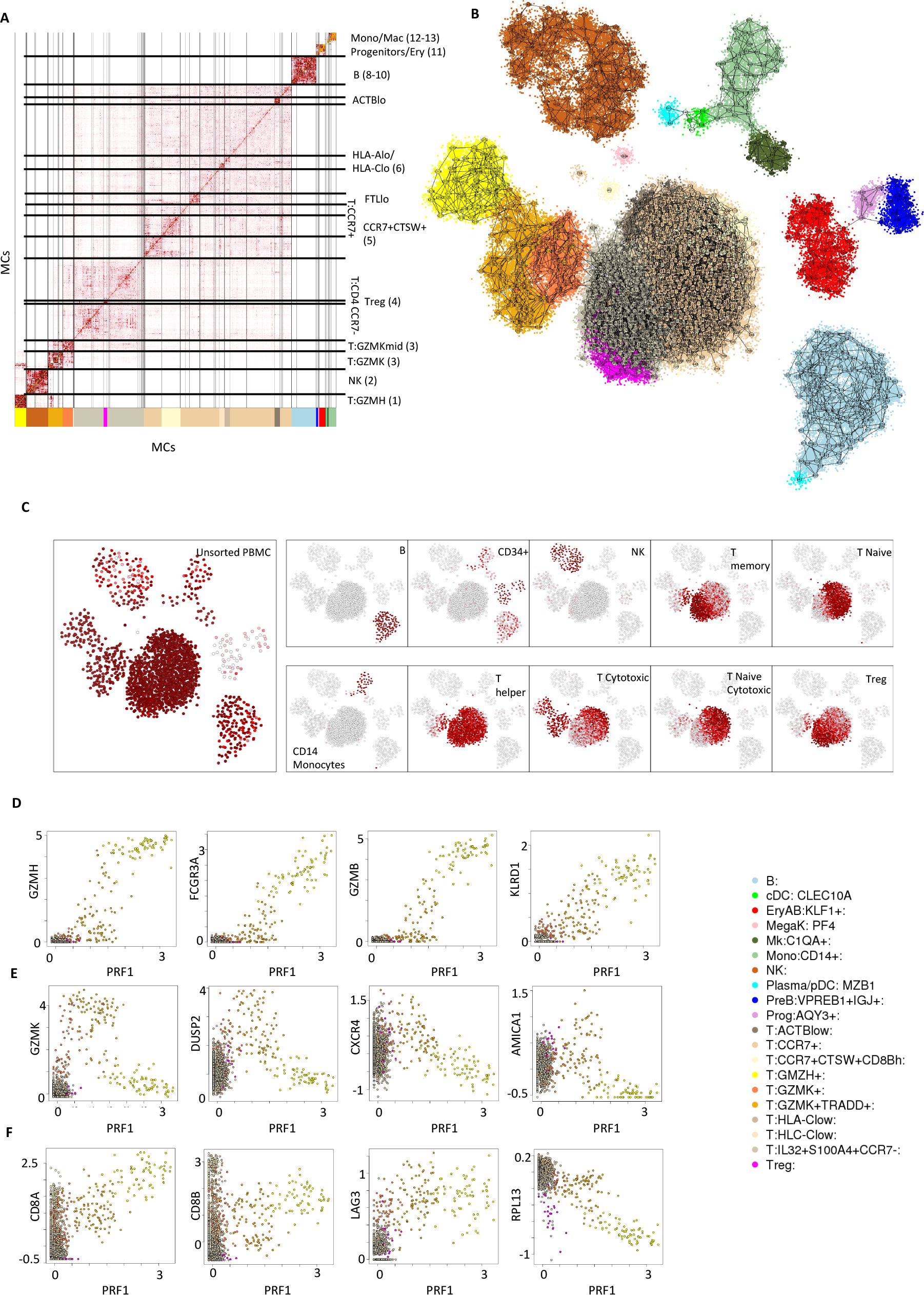
MC analysis of a 160K PBMC multi-batch dataset. **A-B**) Matrix (A) and graph (B) visualization for the similarity structure associating MCs in a model characterizing 162,000 PBMCs. Clusters in the MC matrix are used for linking specific groups of MCs with specific annotation and for color coding. **C**) Shown are the fraction of cells from different sorting batches per MC, color coded white to red to black and visualized using the MC 2D projection as shown in Fig 4B. **D**) Shown are lfp values for MCs in the PBMC 160K model, comparing intensity of Perforin expression (X axis) to several genes correlated with the CD8+ effector program. **E**) Similar to D for genes showing transient activation during the effector program build-up. **F**) Similar to D for CD8 genes, LAG3 (a T-cell exhaustion marker) and a representative ribosomal protein gene.

### Using MCs to define gradients of CD8+ effector T-cell activation

Finally, we demonstrate the potential of applying MetaCell for in-depth analysis of differentiation gradients through analysis of the transcriptional signatures in effector CD8+ T-cells. Activation of the T-cell effector program ultimately depends on expression of units of the cytotoxic granule (granzymes, cathepsins, granulysin) and of the machinery required for perforating target cells (e.g., perforin)^42^. Elevated expression of Perforin 1 (PRF1) is indeed observed in a subset of the CD8+ MCs, spanning a spectrum of intensity from background level to 10-fold enrichment over it. We observed PRF1 enrichment to correlate strongly with multiple additional effector genes, for example granzyme H and B, FCGR3A and KLRD1 (**Fig 5D**), consistent with the idea of a spectrum of transcriptional states with variable effector gene toolkit expression in the blood. Remarkably, we identified a second set of genes showing elevated expression in MCs with low-to-intermediate effector program expression (**Fig 5E**), including most notably granzyme K (GZMK) and the phosphatase DUSP2, but possibly also the chemokine receptor CXCR4 and the adhesion/motility molecule AMICA1/JAML. The effector program expression gradient was also associated with decrease in relative housekeeping gene expression (e.g. ribosomal proteins, **Fig 5F**). We note that the association between the transcriptional gradient of effector genes and temporal or differentiation processes cannot be assumed immediately. It is nevertheless tempting to suggest that effector program activation involves transient expression of the GZMK-linked genes observed here, suggesting several experimental directions for follow up towards a better understanding of T-cell commitment and regulation in the blood and other organs, and in particular within tumors^30,43^.

## DISCUSSION AND CONCLUSION

We introduce here the use of metacells for analyzing scRNA-seq data. Metacells are defined as groups of single cell profiles that ideally represent re-sampling from the same cellular state. In practice, we compute MCs as a graph partition using adequately processed similarities between single cell profiles. We demonstrate that in real data we can construct partitions such that the intra-MC UMI distribution is approximately modeled by the experimental sampling variance, i.e. sparse multinomial sample. We show how to screen for MCs with overdispersion or residual pairwise gene correlations, reflecting deviation from this model and residual intra-MC biological variation. We then demonstrate how the MCs can be used for in-depth exploration of PBMC expression. The analysis methodology we advocate involves direct inspection of the MC adjacency matrix, which provides analysts with complete information about cell type hierarchy and supports clustering at appropriate resolution. Combined with visual examination of correlation patterns between MC-enriched genes, the result is a detailed and unbiased characterization of cell types and expression gradients.

The main property that makes metacells a powerful analysis tool is their ability to increase the signal to noise ratio in the data without introducing biases stemming from mistaken modelling assumptions or over-smoothing of the data. The only manipulation performed by MetaCell on the data is the pooling of highly similar cells, thereby forming a partition of the data. The analyses we present show that, despite enforcing this partitioning, a metacell cover provides accurate local approximations of the expression manifold. At the same time, partitioning entails multiple advantages. Statistically, it greatly reduces the effective number of parameters of the model, making it less prone to over-fitting and to over-smoothing compared with naïve smoothing approaches. For the analyst, it allows for the characterization of well-defined, discrete and highly granular states in a conservative and easy-to-interpret framework.

In cases where residual intra-MC structure is detected in the cover, additional cells can be sampled to refine the MC cover and tighten the approximation. Fundamentally however, in any realistic data set there will always remain some under-sampled behaviors regardless of sampling depth, and our current model will not provide a constructive approach for understanding such behaviors beyond signaling them out as non-homogeneous. Fitting more flexible intra-MC models, capable of accounting for not only sampling noise but also convergent processes such as cell cycle or stress^44,45^, or embedding the metacells in hierarchical or multi-resolution structures^46,47^ should allow for more efficient extraction of the signals of interest. We view the integration of such models as an important future extension of this work.

## METHODS

### Notation and definitions

We assume raw scRNA-seq reads are mapped to genome sequences and assigned to cell barcodes and unique molecular identifiers (UMI) using pipelines that eliminate most of the UMI duplications induced by PCR and sequencing errors. We summarize all UMIs in the molecule count matrix *U* = [*u*_*gi*_] on genes *g* ∈ *G* and cells *i* ∈ *I*. We define *u*_*g*_ as the total molecule count for gene g on the raw count matrix, and *u_i_* as the total number of molecules for a cell (sometime referred to as the cell’s *depth*). The procedures below are designed to robustly define a metacell partition over the cells, which is denoted by a set of cell subsets *M*_*k*_ and a set of outliers *0* such that (⋃_*k*_*M*_*k*_) ∪ *0* = 1. Analysis is performed while filtering outliers and noisy cells, and we therefore do not assume the count matrix to be completely devoid of problematic profiles (e.g. those originating from empty wells or droplets). Nevertheless, we implicitly assume the fraction of completely noisy profiles is relatively low, and some initial threshold of minimal cell UMI count must be employed.

We assume a set of gene features *F* ⊆ *G* is specified and focus our analysis on a similarity graph between cells derived using data from these features (see below). We discuss several strategies for selecting genes in the Appendix. We note that our features represent individual genes rather than principle components or other forms of reduced dimensions. This enables some direct approaches to testing and correcting the gene expression distributions within metacells. It also forces the modelling of similarities and derivation of metacells to work over high dimensional spaces and to account for noise and sparse data directly. Applying the metacell algorithmic pipeline to similarity structures derived using popular dimensionality reduction techniques is easily applicable as well, as we demonstrate in the results section.

### The metacell balanced K-nn cell similarity graph

Grouping of cells into metacells requires modelling of the cell-to-cell similarity in the data. Ideally, this should take into account assumptions on the biological and experimental processes generating the data. However, a well-founded parametric generative model for scRNA-seq data is currently missing, mainly due to the limited understanding of the biological variation in transcriptional states within different cell populations, and the remarkable diversity of coupled (e.g. developmental) and uncoupled (e.g. cell cycle, stress) biological processes that are captured in typical single cell RNA-seq maps. We therefore use a simple heuristic approach for modelling raw pairwise local similarities, which is then refined by additional analysis of the derived cell K-nn similarity structure.

Assume the data is generated by a set of distinct cellular states, where each generates single cell profiles by drawing gene expression independently from log-normal distributions with a fixed variance and gene-specific mean, and then sampling molecules (UMIs) multinomially using the sampled gene concentrations. Given an observed single cell UMI profile *u*_*gi*_, our estimate for the multinomial parameters that generated the profile would be proportional to *u*_*gi*_ + *∊* with some prior parameter *∊*. Moreover, we can approximate the likelihood that two profiles *u*_*gi*_,*u*_*gj*_ were generated from the same log-normal distribution as proportional to ∑_*g*_ (log(*u*_*gj*_ + *∊*) – log(*u*_*gj*_ + *∊*))^2^. Given this rationale, we transform the raw UMI count U on the gene features F as *U′* = [*u*′_*gi*_ = [*log*_2_(*∊* + *u*_*gi*_)]_*g*∈*F*_ and compute the raw similarity matrix using Pearson correlations on the transformed features *R* = [*r*(*u*′_*gi*_, *u*′_*gj*_)]_*ij*_. A simple variation on this procedure may include prior normalization of the U matrix by down-sampling (sampling *min*(*u*_*i*_) UMIs from each cell without replacement) so as to avoid biases associated with improved accuracy (and thereby higher similarity) between deeper UMI profiles. We however avoid down-sampling when the distribution of the number of UMIs per cell is highly variable and correct for the sampling bias when manipulating the similarity graph as described below.

Next, we use the raw similarity matrix *R* to generate a weighted adjacency matrix for a directed cell graph, in which a heavy edge from cell *i* to cell *j* indicates strong attraction of the former to the latter. We first perform a non-parametric transformation by computing *S* = [*s*_*ij*_] = [*rank*_*j*_(*r*_*ij*_)]. Here *rank* is the ranking function, and each row represents the order of similarity between all cells *j* and a specific cell *i*. The *S* matrix is highly non-symmetric, for example when the similarities going from an outlier cell are linking it to members of a large, homogeneous and highly connected cell group. To better control for such effects we perform the following *balancing* operation. We first symmetrize *S* by multiplying ranks *s*_*ij*_ * *s*_*ji*_, followed by initial regularization of edges using a threshold *αK*^2^ on the ranks product:

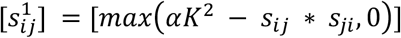

We then perform two rounds of additional regularization, first keeping maximum scoring *βK* incoming edges for each node:

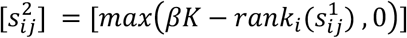

and then further filtering to keep maximum K outgoing edges for each node:

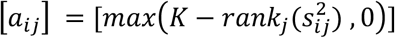

A weighted directed graph G is then constructed using [*a*_*ij*_] as the weighted adjacency matrix. Note that nodes with degrees lower than K are possible following this procedure, since outlier cells may become disconnected or poorly connected during the balancing operations.

### Seeding and optimizing graph partitions

We now describe how the MetaCell partition is produced, i.e. how the graph is partitioned into similarly-sized, homogenous subgraphs, to be later used for quantifying expression states. Since these subgraphs are not necessarily maximal, a single densely-sampled population may be represented by multiple MCs. The optimal partition should be as granular as possible while still allowing for robust statistical characterization of the underlying expression profile. In the current procedure we settle for obtaining similarly-sized subgraphs that pass some minimal size threshold. A major challenge is to effectively covering cell cohorts that are often non-uniformly sampled, consisting of dense populations as well as small cell groups representing rare states. The preliminary seeding stage addresses this challenge by making sure that all cells stand a good chance of having a subgraph formed in their immediate neighborhood.

We partition the balanced similarity graph G into dense subgraphs using an adaptation of k-means to graphs. Let the parameter K define the typical desired size of subgraphs in the partition (which is also the maximum outdegree of the graph G as constructed). Denote by *N*^*out*^(*i*) the set of graphic outgoing neighbors of *i*. We initialize an empty assignment of cells to subgraphs *mc*(*i*) = –1, define the set of covered nodes as *C* = {*i* | *mc*(*i*) > –1} and the *cover-free* score for each node as *f*(*i*) = |*N*^*out*^(*i*)- *C*|. We then sample subgraph seeds using an iterative procedure:

Initialize *k* = 0

While 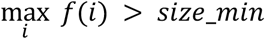 do:

> sample a new seed cell *j* by drawing a sample from cells in *I* – *C* with weights proportional to *f*(*i*)^3^ update *mc*(*u*) = *k for u* = *j*,*u* ∈ *N*^*out*^(*j*) – *C*
>
> Increment k and Update *C, f*

We terminate seeding using a minimum subgraph size parameter *size_min* < *K*. When we meet the stop criterion, cells that are not associated with a seed (i.e. cells for which *mc*(*i*) = –1) have at most *size_min* uncovered neighbors, and in particular will almost always have at least one covered neighbor (since the degree in the balanced graph is typically K).

The seeding step produces an initial set of subgraphs *M*_*k*_ = {*i*|*mc*(*i*) = *k*} that forms a basis for further optimization. Define the outgoing association of each cell to a subgraph as 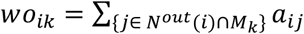 (recall *a* are the graph weights), and analogously the incoming subgraph association for each cell as 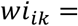 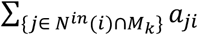. The combined cell-to-subgraph association is computed by multiplying the outgoing and incoming weights and normalizing by the respective subgraph size: *w*_*ik*_ = *wi*_*ik*_ *wo*_*ik*_/|*M*_*k*_|^2^. We use this scoring scheme to iteratively optimize. as well as complete the initial graph cover, so that it includes all cells:

Until convergence:

> Select a cell *i*
>
> Reassign *mc*(*i*) = *argmax*_*k*_ *w*_*ik*_
>
> Update weights

Convergence is defined by deriving a partition in which all cells are associated with their highest scoring subgraph. To enforce convergence (which is not guaranteed to occur in general) we slowly increase the score association between cells and their current subgraph after each reassignment. This is especially useful when a large subset of cells (i.e. larger than K) are very homogeneous, which may result in unstable exchange of nodes between several modules covering this subset.

After convergence, there are no formal guarantees on size distribution of the subgraphs produced by the algorithm. Empirically, however, the connectivity of the graph (maximum K outgoing edges) and the seeding process promote a relatively uniform cover partition prevent convergence toward solutions with very large subgraphs. Rare cases of cells that reside in connected components whose size is smaller than *size_min* and were left uncovered during seeding are defined as outliers.

Importantly, the complexity of the entire procedure (seeding and optimization) is linear in the number of cells and the maximum degree K (or alternatively, linear in the number of edges in the graph). An efficient implementation of the algorithm therefore scales well to large datasets, as does its integration within an extensive resampling strategy, as we discuss next.

### Resampling graph partitions and computing metacells

We improve the robustness of the above randomized graph partition algorithm using a resampling approach. Given the balanced graph G, we generate a series of subgraphs *b* = 1..*N*_*B*_ (typically *N*_*B*_ = 500) by sampling cells independently without replacement with probability *ρ* (typically *ρ* = 0.75) and adding all edges connecting them, forming *G*^*b*^ = (*V*^*b*^, *E*^*b*^), *V*^*b*^ ⊂ *V*, *E*^*b*^ ⊂ *E*. For each resampled *G^b^* we apply the partition algorithm, thereby generating a set of partial graph partitions *mc*^*b*^(*i*) *for each i* ∈ *V*^*b*^. We summarize all partitions using the matrices *O* = [*o*_*ij*_] and *C* = [*c*_*ij*_] specifying how many times the pair of cells *i,j* were resampled together, and how many times they were both assigned to the same subgraph in the resampled partition, respectively. We then define the resampled cooccurrence matrix as 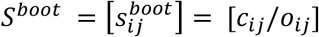.

The values in *S*^*boot*^ are now used to compute a weighted, non-directed graph, discarding the original correlation distances. We compute for each cell *i* the value of the *K*^*boot*^ highest frequency neighbors (denoted *T*_*i*_), and then define a co-occurrence threshold for each pair of cells using the maximal of the two critical values multiplied by a factor *T*_*ij*_ = *max*(*T*_*i*_, *T*_*j*_) * 0.5. Pairs with 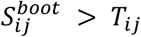 are used as the edges in a new graph denoted as *G*^*boot*^ on all cells. Note that *G*^*boot*^ is still non-symmetric, as setting fixed thresholds on edges implies that nodes in large and diffused clusters will have a lower *T_i_* values and thereby higher degree than nodes in tight and robust clusters that always cluster in the same subgraphs. The parameter *K*^*boot*^ provides users of the algorithm with flexible control over the degrees in the derived graph. The final partition solution is obtained by re-applying the same partition algorithm on the graph *G*^*boot*^, resulting in a new set of subgraphs *M*_*i*_ and a potential list of outliers. This solution is subject to further filtering and verification, as described next.

### Filtering clear parametric outliers from a metacell cover

As commented above, even though we lack a proper parametric model for single cell RNA-seq, our idealized metacell cover is expected to group together single cell profiles that are consistent with multinomial sampling, with parametrization allowing for some overdispersion. Testing a given metacell cover for gross inconsistencies with this assumption can help detecting outlier cells emerging from experimental errors (such as doublets), as well as diagnose rare states that are not sufficiently abundant to define a separate metacell. We currently approach this detection problem heuristically, by summarizing the metacell’s pool frequencies:

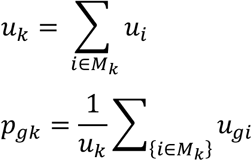

and computing an approximate, regularized observed/expected value for each gene and cell:

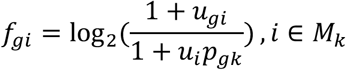

Note that the regularization (adding 1 to observed and expected count) implies that high fold change values cannot be attained for genes with very low overall UMI counts. However, this regularization is sufficient to ensure robust detection of clear outliers. Cells with one or more genes showing high *f*_*gi*_ values are labeled as potential outliers and removed from their metacell cover prior to in-depth quantitative analysis of the model.

### Verifying metacells homogeneity

Outlier filtering does not guarantee metacell homogeneity in cases where two distinct and significantly separated transcriptional states are grouped together. To screen for such scenarios, we attempt to cluster cells within each metacell *M*_*k*_ de-novo. Clustering is performing by applying the DBSCAN density-based clustering algorithm to the intra-metacell similarity matrix, computed as the correlation distances described above but restricted to genes exhibiting mildly high intra-metacell variance. If more than one cluster is detected, we split the metacell accordingly. In practice, metacells almost never include hidden sub-clusters and testing for splits is used mostly for validation purposes.

### Defining the metacell gene expression profile

We approximate the gene expression intensity within each metacell by a regularized geometric mean:

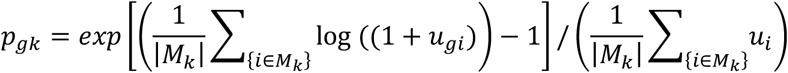

We then quantify relative expression as the log fold enrichment over the median metacell value:

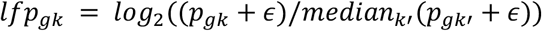

Note that the *lfp* values are affected by the composition of metacells in the dataset up to a constant, and that *∊* (typically set to 10^−4^) should be adapted to the typical total molecule count within a metacell.

### Metacell regularized force-directed 2D projection

Projection of single cell RNA-seq data onto a twodimensional plane provides analysts with a compact and intuitive visual encoding of the similarity structure underlying the data, and is one of the most popular tools practiced by scRNA-seq analysis. A 2D projection must be used cautiously, since it is difficult and often infeasible for such a minimal representation to support differentiation trajectories or quantitative spectra of transcriptional states. We use the Metacell cover to regularize the similarity graph among single cells and therefore simplify their projection as follows. We start by projecting edges in the graph G over metacells:

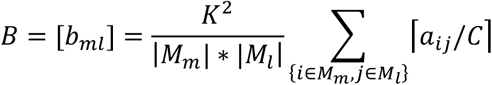

(here *C* = *median*_*k*_(|*M*_*k*_|) is a scaling constant). We symmetrize B by replacing it with B’, the sum of its row and column-normalized forms, and retain as candidate edges only pairs for which *b*′_*ml*_ > *T*_*edge*_. We then construct a graph over the metacells *G*^*M*^ = (*M*, *E*^*M*^), by adding the *D* highest scoring candidate edges (if they exist) for each metacell. This results in a graph with maximum degree *D* and any number of connected components. We compute coordinates (*xm*_*k*_, *ym*_*k*_) for each metacell by applying a standard force-directed layout algorithm to the graph *G*^*M*^. We then position cells by averaging the metacell coordinates of their neighbor cells in the original balanced graph *G*, but filter neighbors that define a metacell pair that is not connected in the graph *G*^*M*^. Averaging allows for layout flexibility along one or few edges in the metacell graph when positioning large cell clusters that are dissected by several metacells.

### Implementation

We implemented MetaCell using a combination of C++ and R code and parallelization over multi-core machines. On a strong Xeon-E5-2660 dual-CPU machine the entire analysis pipeline for a small 8,200 cells dataset, including bootstrap iterations and computing 2D visualizations, required 2 minutes and a maximum of 4.8GB of RAM. The entire analysis pipeline for a 160K cells dataset required 112 minutes and a maximum of 79GB RAM on the same machine.

### Evaluating within-MC homogeneity

Following the computation of the MetaCell partition, our pipeline produces diagnostic statistics and plots to evaluate the level of adherence of the metacells to a multinomial sampling model. To visualize large-scale adherence across all genes, we produce per MC plots comparing the coefficient of variation and the fraction of zero counts to the expected under a Poisson model (see examples in **Fig S5**). In addition, we visualize adherence to binomial sampling of the top enriched genes per MC by plotting the observed distribution of UMI count and the same distribution sampled from a binomial model (see examples in **Fig 2D**). For both observed and expected, counting is done after down-sampling all cells within a metacell to uniform total counts. Finally, global diagnostic matrices over all MCs and marker genes (see example in **Fig 2E**) are computed as follows: We down-sample the UMIs to uniform total counts per MC, and compute the binomial likelihood of the observed counts, as well as their over-dispersion (observed divided by expected variance). We average these statistics over multiple down-samples, and repeat the whole procedure over 999 fake count matrices drawn from the per-MC multinomial model. Per gene and per MC, we compute the empirical p-value of its likelihood with respect to the binomial null. We output the p-values and the overdispersion values, and visualize a summarizing heatmap of the latter. Note that when computing binomial statistics we down-sample with respect to feature and enriched genes only, and that the expected distributions are derived from the pool frequencies constrained to these genes.

### Comparing local approximation accuracy using expression prediction

We designed a cross-validation experiment to quantify how well the MetaCell partition captures local cell-to-cell similarities. We divided the gene set into 100 folds, and leaving out each fold at a time computed cell-to-cell similarities on the remaining genes using four different strategies. We next used these similarities to predict, per cell, the expression level of the left-out genes. Finally, we compared the quality of predictions across all genes. A model that captures accurately local similarities in the expression manifold is expected to produce accurate predictions.

The compared approaches are: 1) Predicting using the per-metacell pool frequencies. 2) Predicting using the pool frequencies among the top 50 neighbors according to the raw MC similarity matrix *R*. 3) Predicting using the pool frequencies of the top 50 neighbors according to Euclidean distances in Seurat’s PCA space. 4) Predicting using the weighted pool frequencies of all cells, where the weights are set as MAGIC’s diffusion similarities (more specifically, MAGIC’s powered Markov affinity matrix). Pool frequencies were computed as regularized geometric means, denoting by *w*_*i*_ the weight of cell *i* in the pool (for strategies 1-3 all weights are 1):

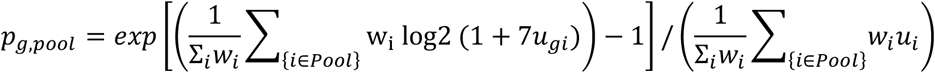

The extent of over-fitting was tested by avoiding the cross-validation design and computing a single similarity matrix using all genes per modeling approach. Regardless of whether cross-validation was used, a cell was never a part of its own prediction pool when comparing prediction accuracy (Fig 3B, 3C). In contrast, for plotting the gradients (Fig 3D, 3E) the predicted values were generated using all genes and all cells, as in a typical analysis.

Combining Seurat and MetaCell’s filtering criteria, only cells with at least 800 UMIs, number of expressed genes between 800 and 4,000, and mitochondrial gene fraction below 0.1 are included. We omitted from the modeling and the evaluation mitochondrial genes and immunoglobulin genes. For MetaCell we used MC size parameter K=100 and 500 down-samples of 0.75 of the data during the graph resampling stage. For Seurat(package downloaded on 18/3/26) we used gene selection parameters x.low.cutoff = 0, y.cutoff = 0.8, negative binomial scaling over mitochondrial fraction and number of UMIs, and 40 PCs. For MAGIC (code downloaded on 18/3/19) we used 30 PCs, k=5, ka=4, epsilon=1 and t=6.

### Whole organism scRNA-seq analysis

For the *Caenorhabditis elegans* map, we analyzed the whole-organism single-cell dataset published by Cao et al.^39^ and generated using methanol-fixed larval L2 stage cells and a split&pool scRNA-seq strategy. We started from a UMI matrix containing 41,449 single cells. We filtered out cells with less than 100 and more than 8,000 total UMIs. We used MetaCell to select marker genes with the following criteria: (1) a normalized size correlation below −0.1 and/or a niche score over 0.1, (2) a minimum of 300 total UMIs observed, and (3) a minimum of 3 UMIs observed in at least three single cells. For MetaCell, we used MC size parameter K=150 and 1,000 down-samples of 0.75 of the data during the graph resampling stage. We computed the final partition from the coclustering matrix using a size parameter K=30, a minimum MC size parameter of 30 and alpha=2. We filtered outlier cells using a filtering parameter T_lfc=4, resulting in a final filtered set of 38,149 cells.

For *Schmidtea mediterranea*, we analyzed the whole-adult single-cell dataset published by Fincher et al.^40^ and generated using fresh cells from whole-adult and head area planarian samples and the Drop-seq scRNAseq technology. We started from a UMI matrix containing 58,328 single cells. We filtered out cells with less than 500 and more than 18,000 total UMIs. We used MetaCell to select marker genes with the following criteria: (1) a normalized size correlation below −0.1 and/or a niche score over 0.05, (2) a minimum of 300 total UMIs observed, and (3) a minimum of 3 UMIs observed in at least three single cells. In the graph partitioning stage we used the same parameters as in the *C. elegans* analysis. We filtered outlier cells using a filtering parameter T_lfc=4.5, resulting in a final filtered set of 56,627 cells.

### Fine clustering using Seurat

Seurat’s clustering algorithm was used for producing a high-resolution clustering of the 160K PBMCs dataset by applying the following procedure: Data was log-normalized and scaled to 10,000 UMIs per cell, 1,000 genes with top variance/mean ratio were used as highly variable genes, these genes were rescaled by regressing on per-cell number of UMIs, and PCA reduction to 45 dimensions was applied to the rescaled variable genes. In order to generate a fine clustering solution we set Seurat’s resolution parameter to 100, using the approximation parameters nn.eps=0.5 and n.start=10, which yielded 817 clusters. We note that Seurat is typically executed with much lower resolution values (0.6-3).

## DECLARATIONS

### Software and data availability

MetaCell’s open-source code is maintained on Bitbucket and available from https://bitbucket.org/tanaylab/metacell/src/default/

The PBMC data sets were downloaded from the 10x Genomics website:
8K unsorted - https://support.10xgenomics.com/sinhgle-cell-gene-expression/datasets/2.1.0/pbmc8k

68K unsorted - https://support.10xgenomics.com/single-cell-gene-expression/datasets/1.1.0/fresh_68kpbmc_donor_a

10 bead-enriched subpopulations - https://support.10xgenomics.com/single-cell-gene-expression/datasets/1.1.0/
(cd14_monocytes,b_cells,cd34,cd4_t_helper,regulatory_t,naive_t,memory_t,cd56_nk,cytotoxic_t,naive_cytotoxic)

C.elegans L2 larva stage dataset was obtained from http://atlas.gs.washington.edu/worm-rna/data/

Planaria whole-organism dataset was obtained from https://www.ncbi.nlm.nih.gov/geo/query/acc.cgi?acc=GSE111764

## Acknowledgements

We are grateful to all members of the Tanay group for discussion.

---

**Supplementary Figure 1:**
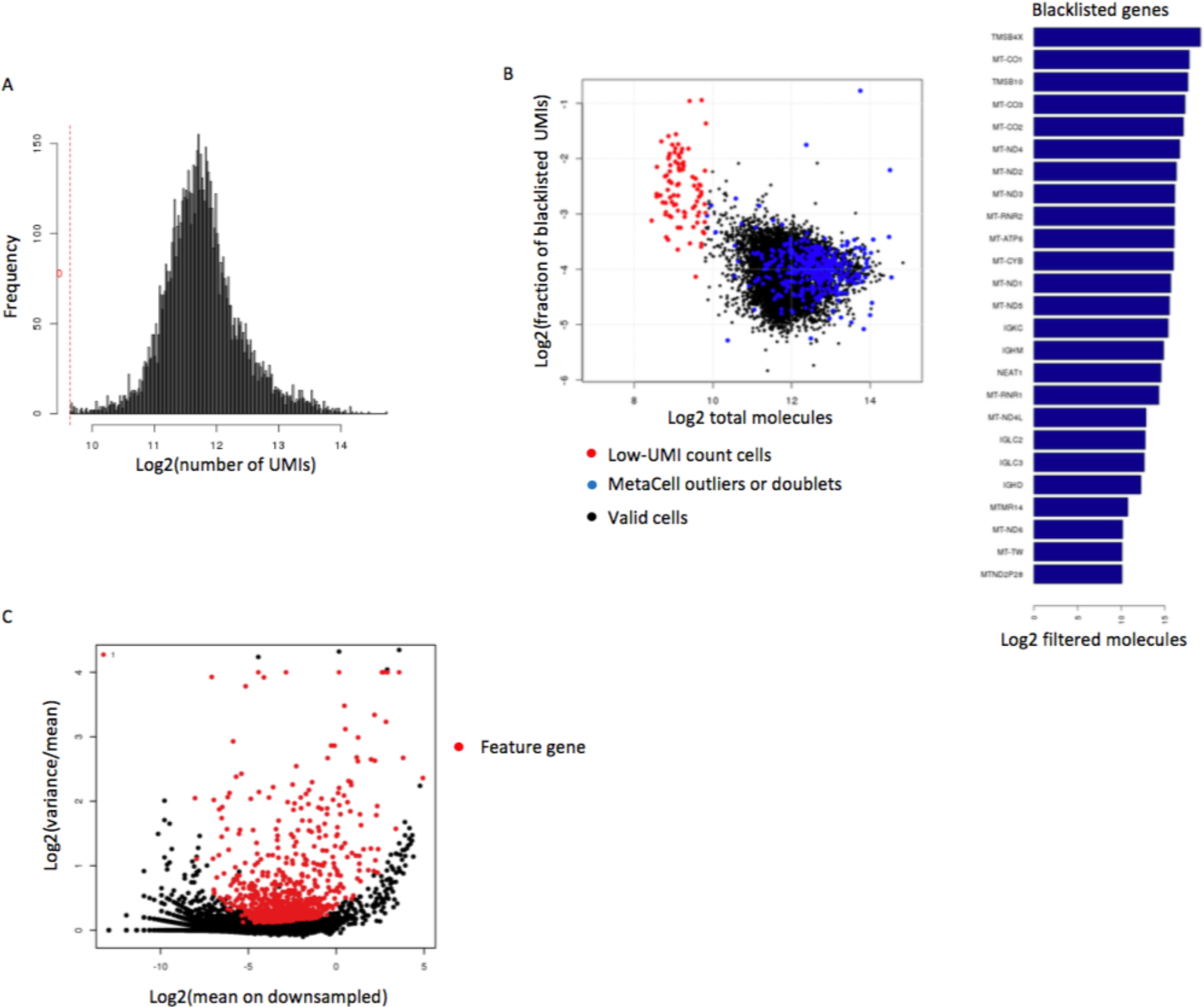
PBMC filtering and feature gene selection. **A**) distribution of number of UMIs per cell in the PBMC 8K dataset, after filtering cells with less than 800 UMIs. **B**) Total number of UMIs (X axis) vs total number of UMIs from genes marked as blacklisted (Y axis). Cells that were filtered based on low UMI count are shown in red. Cells filtered by the MetaCell pipeline based on outlier gene detection or doublet MC annotation are shown in blue. Total number of filtered UMIs for the most abundant filtered genes is shown in bars at the right. **C**) Gene’s mean UMI count vs normalized variance across the entire PMBC 8K dataset. All statistics are computed on down-sampled UMI matrices. Genes selected as features for computing cell-cell similarity are marked in red.

**Supplementary Figure 2:**
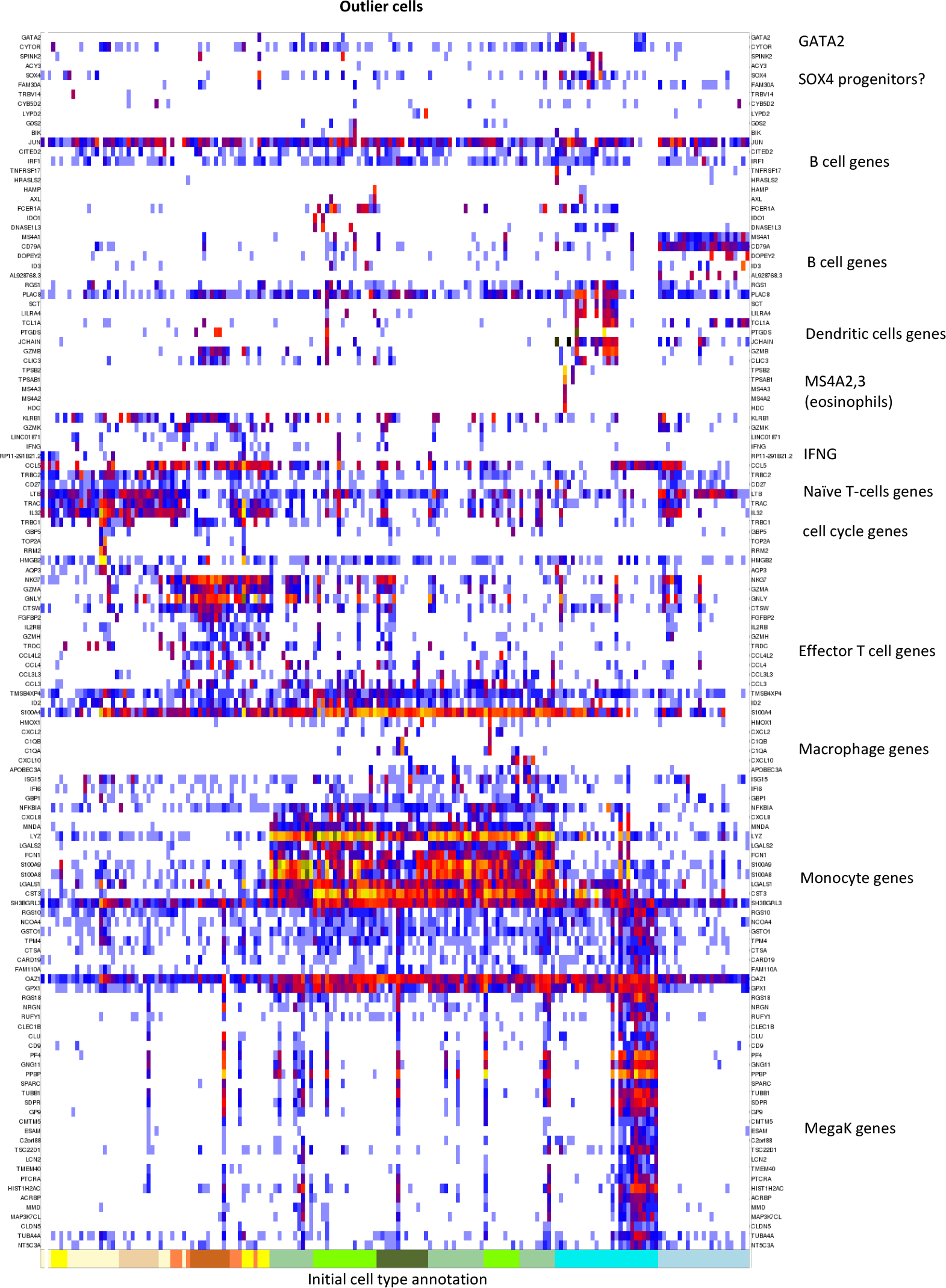
Outlier matrix (enlarged version of Fig 1B). Color coded UMI counts for outlier genes (rows) and cells (columns) are shown. Note that genes that define outlier behavior for one cell are frequently valid marker genes for other cells, in particular when outlier cells represent a doublet behavior.

**Supplementary Figure 3:**
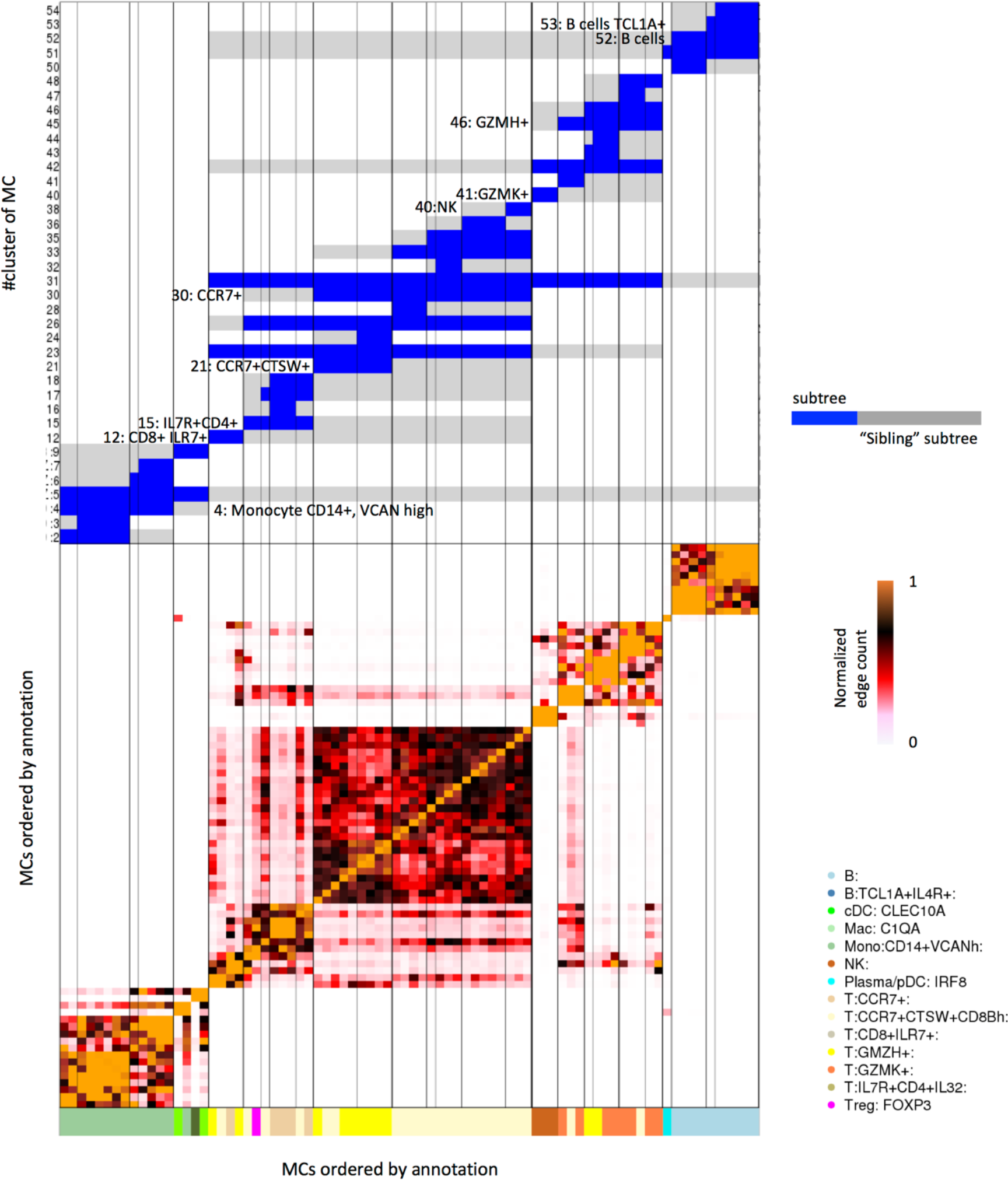
Annotating the PBMC 8K MetaCell model using a clustered MC adjacency matrix (color coded heat map, lower part). Identification of subtrees in a hierarchical MC clustering (using a standard Ward R implementation) based on the adjacency matrix (upper panel, subtrees marked in blue, sibling subtree marked in gray), is then followed by detection of enriched genes per subtree. This facilitate the labeling of groups of metacells to specific known or putative biological functions. Labels next to subtree bars indicate the subtree number and its annotation. Note that the granularity of annotation greatly depends on analyst decisions and goals.

**Supplementary Figure 4:**
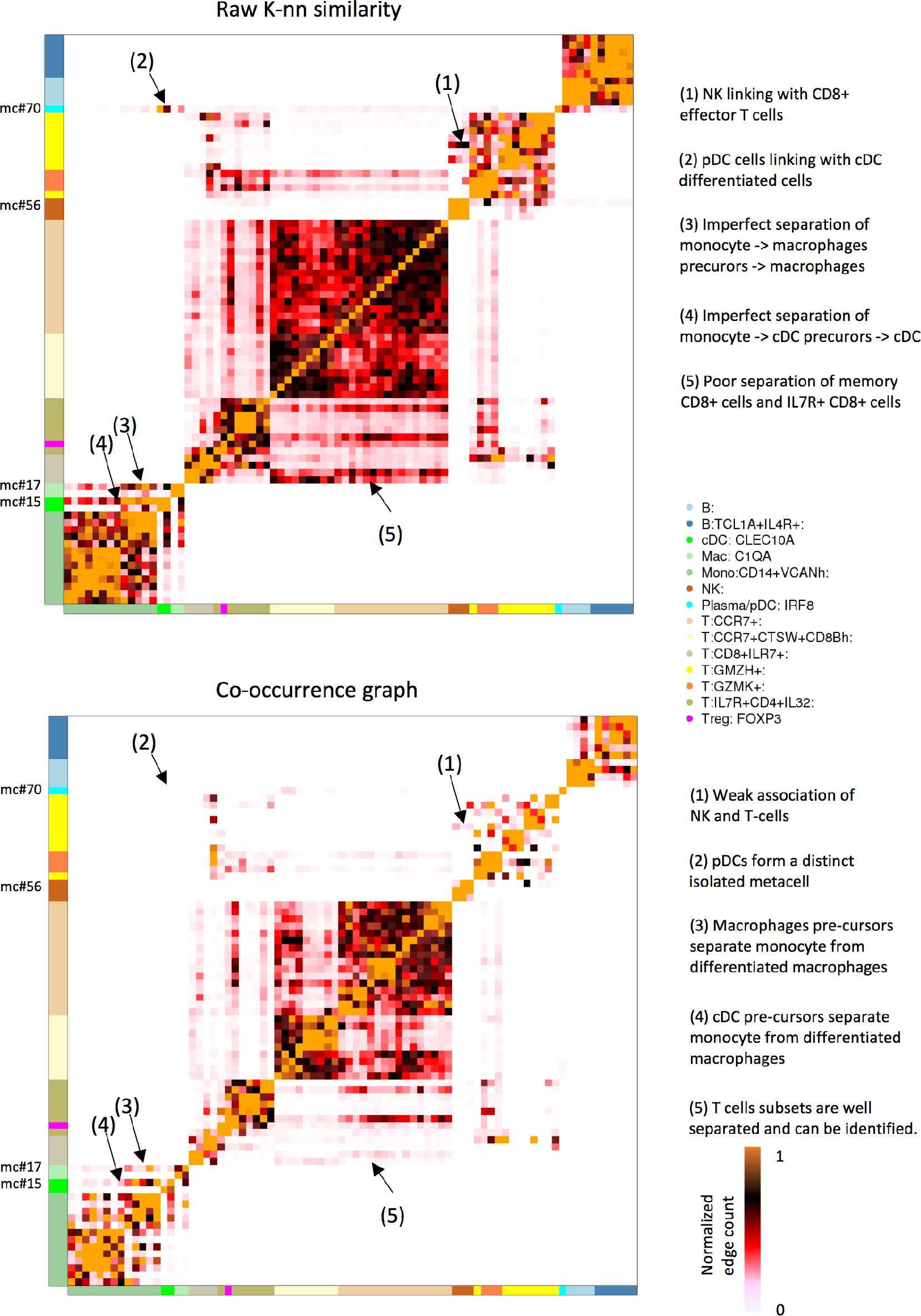
Changes in MC adjacencies during graph balancing. Shown are color-coded MC adjacency matrices for the PBMC 8K model using the raw K-nn graph and the co-occurrence graph (matching with Fig 2B). Specific adjacencies are highlighted (numbered 1-5) to exemplify some key effects of the graph balancing procedure.

**Supplementary Figure 5:**
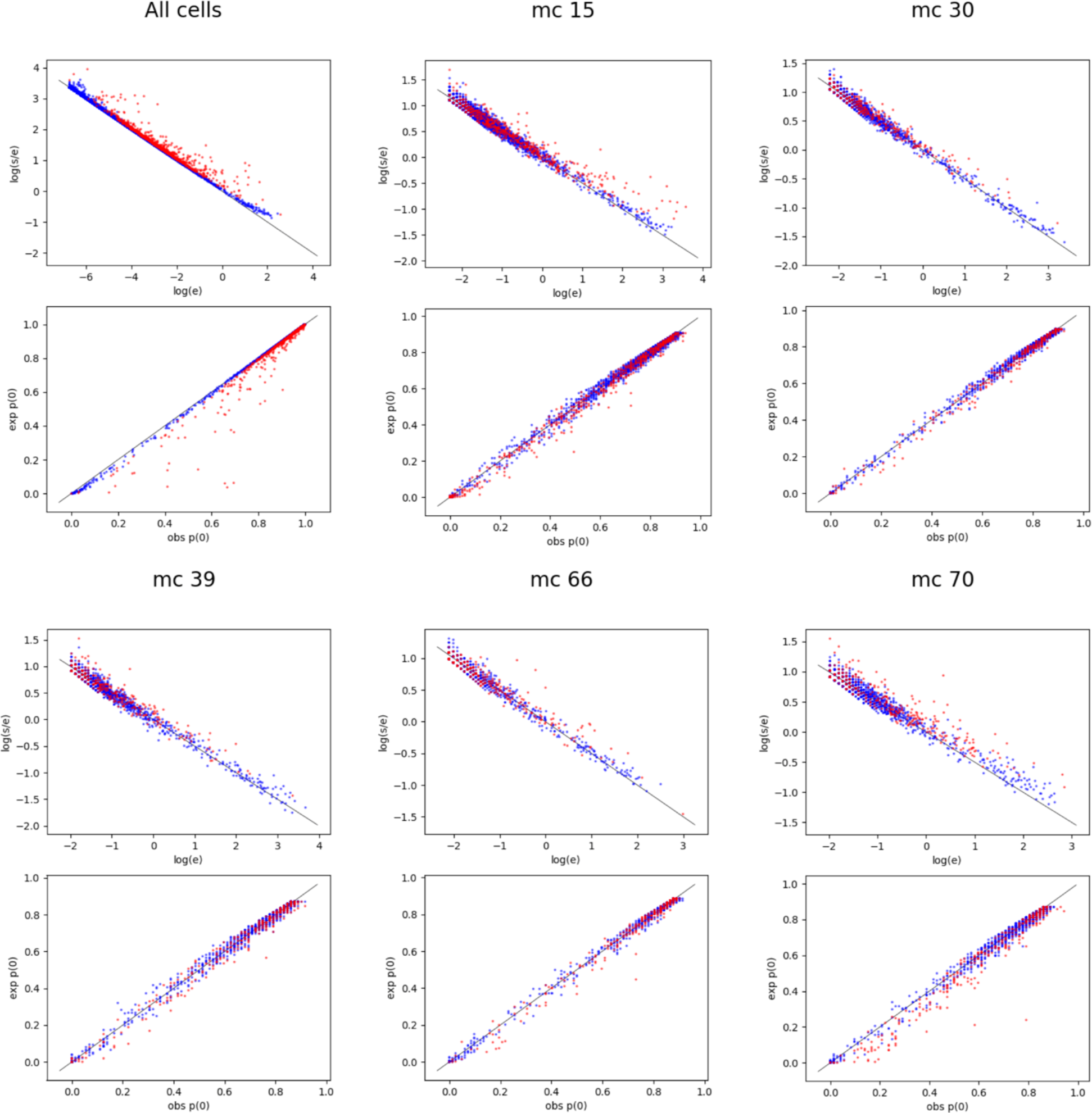
Poisson plots evaluate over-dispersion within MCs in the 8K PBMC data set. Per MC, we down-sample all cells to uniform depth, and compute per gene the mean e and standard deviation s of expression values. We plot log(s/e) vs log(e), which should yield a constant −0.5 slope for perfect Poisson approximation. We also plot the expected vs observed fraction of cells with 0 UMIs. Only genes with at least 10 down-sampled UMI’s per MC are shown, and feature genes are colored red. For MCs whose residual variance highly exceeds multinomial sampling variance (see for example MC #70) many genes show overdispersion and zero inflation.

**Supplementary Figure 6:**
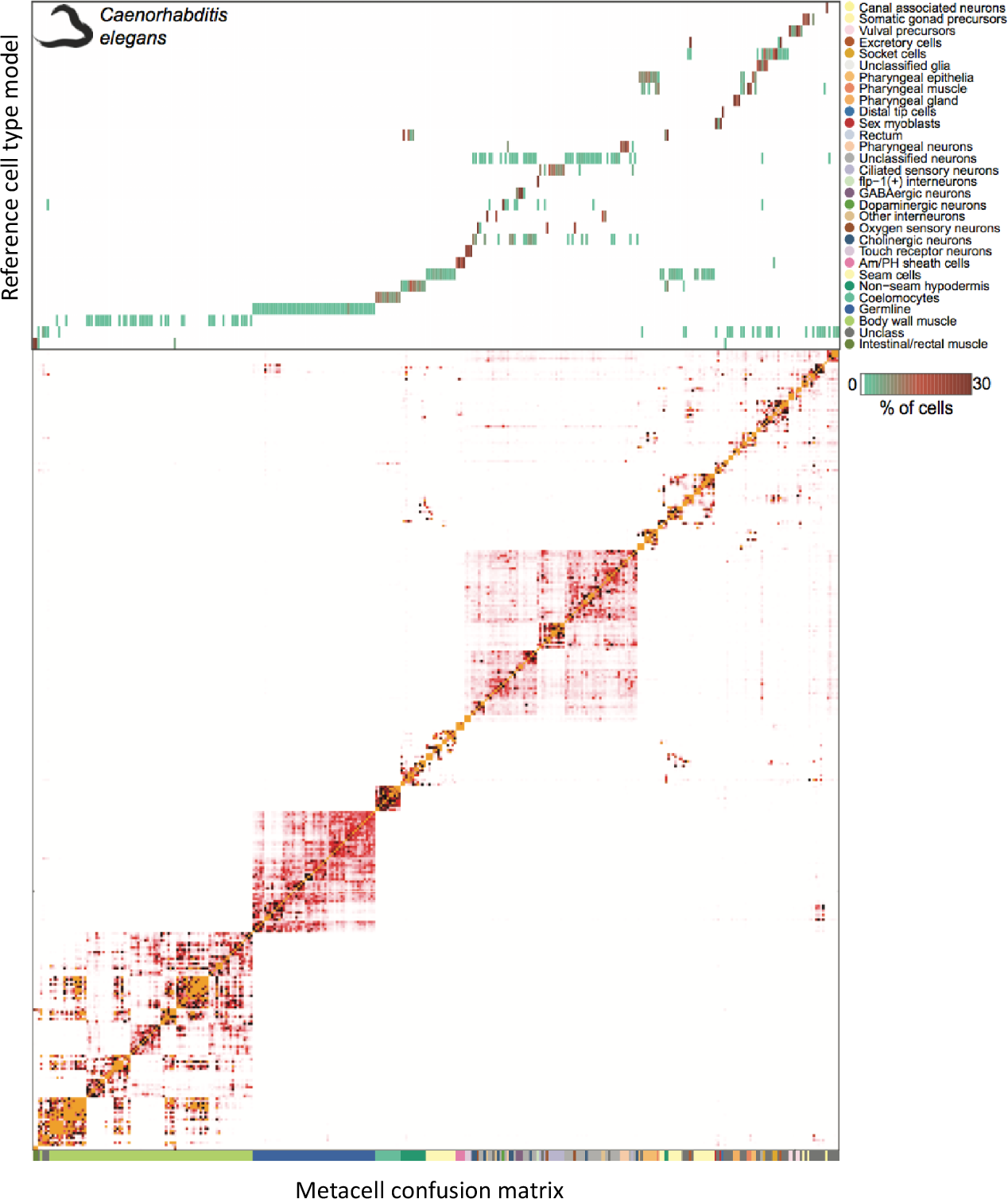
Comparison of the *C. elegans* L2 larva MC model versus the reference cell type model. Top: heatmap showing, for each original cell type in Cao et al. (rows), the distribution of single cells across the metacell (columns). Bottom: heatmap representing the similarity structure between metacells, based on the number of edges in the balanced MC graph that links two cells associated with different MCs. Metacells are color-coded as in Figure 4.

**Supplementary Figure 7:**
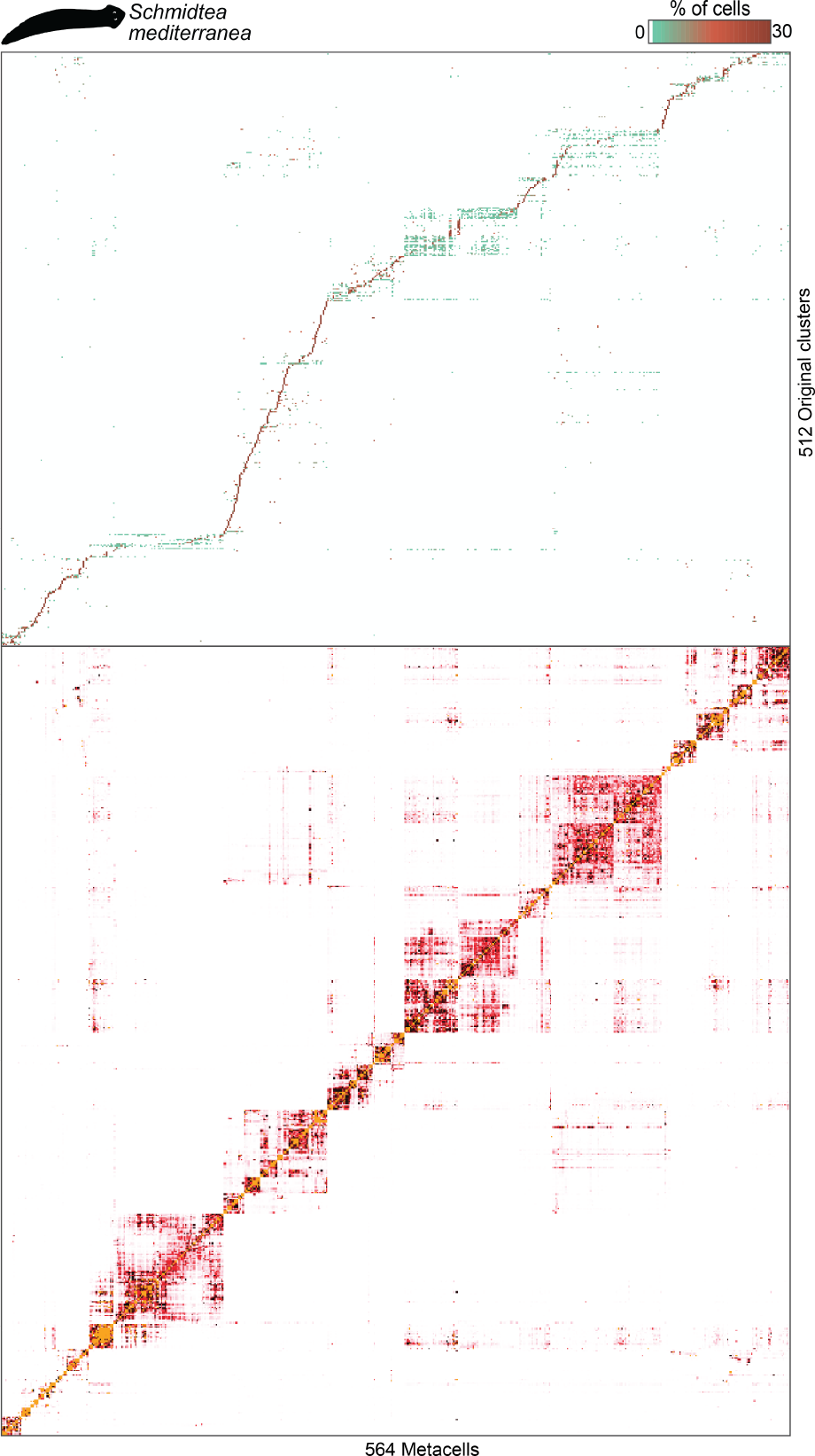
Comparison of the *S. mediterranea* whole-adult MC model versus the original clustering model. Top: heatmap showing, for each original cell cluster in Fincher at al. (rows), the distribution of single cells across metacells (columns). Bottom: heatmap representing the similarity structure between metacells, based on the number of edges in the balanced MC graph that links two cells associated with different MCs.

**Supplementary Figure 8:**
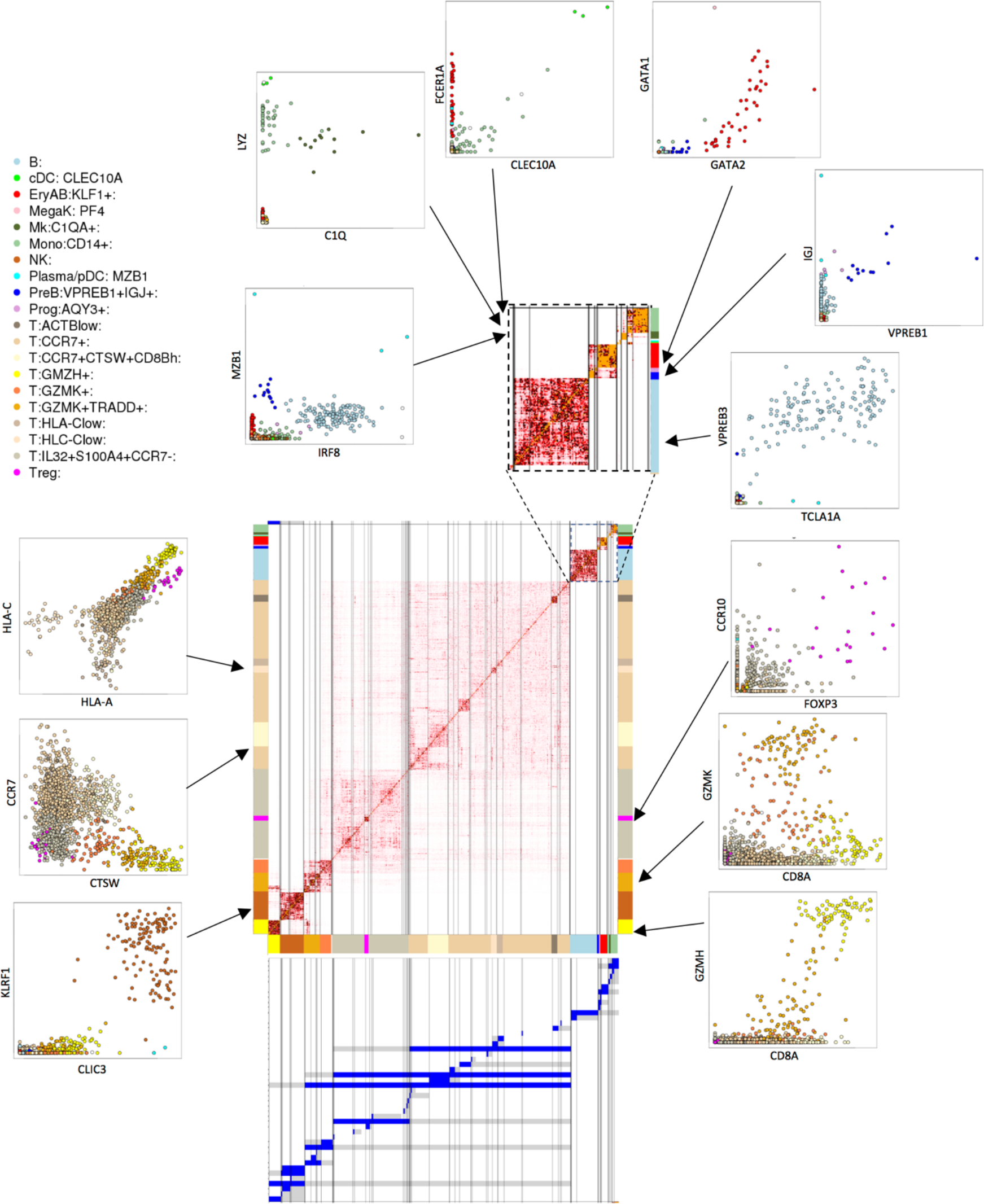
Supporting annotation of MC groups in the PBMC 160K model. The MC adjacency matrix (center, color-coded as described above) defines a hierarchy of clustered MCs (depicted as blue/gray bands, bottom: subtrees marked in blue, sibling subtree marked in gray). Specific clusters of MCs are then annotated based on their gene expression signatures. Shown here are select comparisons of gene enrichment (lfp values in X and Y), supporting some key annotations in the PBMC model. Adjacencies between myeloid, progenitors and B cell MCs are enlarged (top right) for clarity.

**Supplementary Figure 9:**
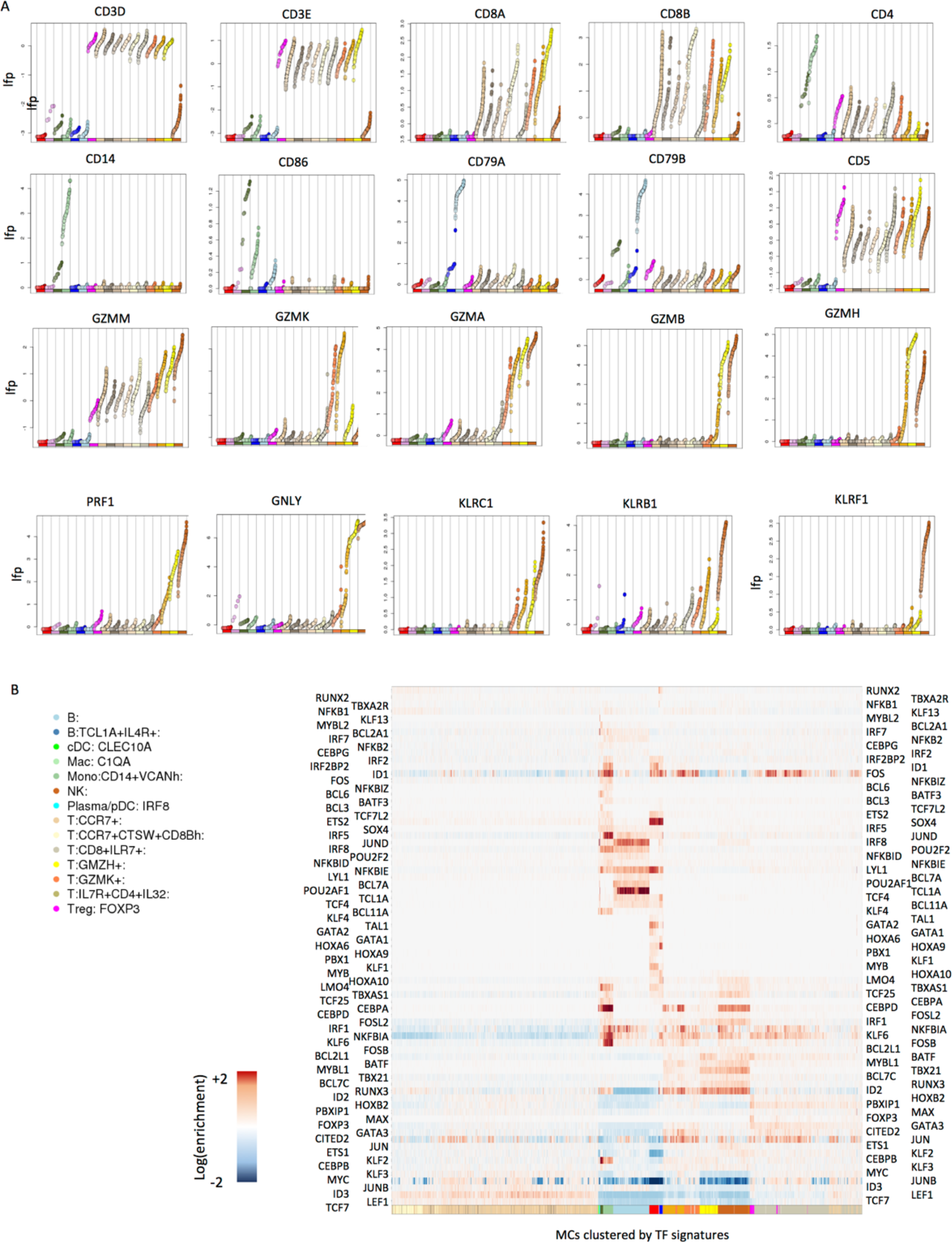
Effector genes are expressed in a convergent fashion. **A**) Shown are MC enrichment values for select surface markers and effector genes, ordered according to MC cell type annotation. Note the expression of granzyme and killer genes across multiple subsets of T-cells and NK cells. **B**) Reference mapping of transcription factor expression in the PBMC 160K model. Abundant groups of (mostly naive) T-cells are defined by relatively few enriched TFs (TCF7, LEF1). Other TFs (e.g. the early-immediate regulators JUN and FOS, or CEBPD) are enriched in multiple cell types. Relatively few TFs are precisely restricted to highly specialized programs (e.g. TCL1A, SOX4).

**Supplementary Figure 10:**
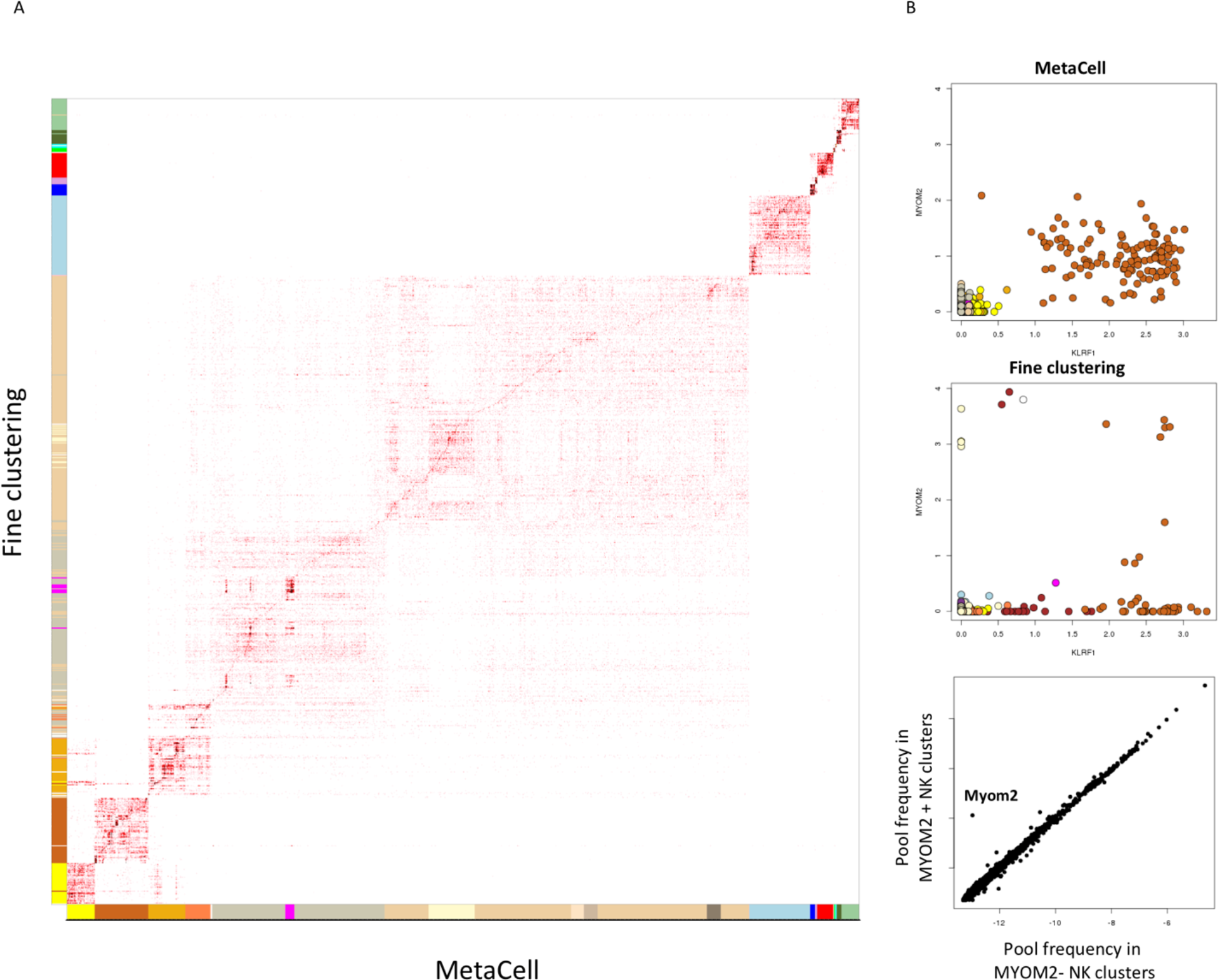
Comparing MetaCell to fine clustering. **A**) The matrix shows the number of cells within each MC (columns) and cluster (rows), where clusters are derived by applying Seurat with parameters forcing high resolution clustering. **B**) Shown are log fold change enrichment per MC (top) and cluster (middle) for KLRF1 (an NK cell marker) and the MYOM2 gene. MYOM2 is generally enriched in NK MCs, but shows highly specific enrichment in specific NK clusters in the fine clustering solution. Bottom: Specific MYOM2 enrichment is suggested to represent overfitting by analysis of overall gene expression in cells within NK clusters with high (Y axis) or low (X axis) MYOM2 levels. Each marker is a gene, and the plot demonstrates no additional genes to be co-variating with MYOM2 within NK cells.

## Appendix: Two strategies for selection of feature genes

Selecting genes *F* for initiating the modeling of cell-to-cell similarities can be done based on a standard identification of high variance genes, or, in cases where normalizing cells to uniform depth poses a problem, it can be achieved through analysis of the correlation of putative feature genes with cell depth.

To select high variance genes, we define a UMI count threshold *T*_*u*_ = *quanile*_*i*_(*u*_*i*_, 0.1) and create a matrix *W* = [*w*_*gi*_] by down-sampling *T*_*u*_ molecules from each cells for which *u*_*i*_ > *T*_*u*_, while discarding all other cells. Rows in the matrix W are defined by their mean UMI count, 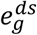, and by their variance 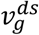. The variance should be affected by three components. First, since molecules are being sampled from each cell, the sampling variance is expected to be in the order of the mean number of sampled molecules, with a distribution that follows a binomial model (*sampling variance*). Second, RNA concentrations of a gene within homogeneous cell populations are subject to stochastic control that contributes additional variance to our sample (*stochastic variance*). Third, when observing heterogeneous single cell populations, genes that are differentially expressed will be affected by additional variance associated with the sampled sub-populations or cell types (*regulatory variance*). The variance in low expression genes 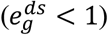 will be dominated by the sampling component, while for genes with higher variance it is difficult to separate stochastic from regulatory variance. We detect genes whose variance seems more than stochastic as those whose regularized variance 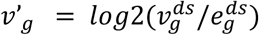 is high given their mean expression. Specifically, we compute the empirical trend *ν*′(*e*) as a function of 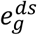 using a moving median, and recalibrate the variance over this trend as 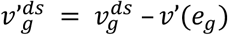. Genes with max 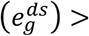 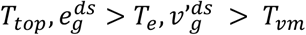 are selected as features.

Alternatively, when cell depth is highly variable we may prefer to avoid down-sampling the matrix and instead directly correct for the effect of this variation. For each gene *g*, denote by 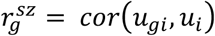 its Pearson correlation with the cell depth 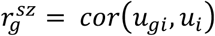. This statistic correlates strongly with mean expression, as increased expression implies lower sampling variance and hence increased correlation with cell depth. For a given level of mean expression this correlation reflects the fraction of sampling variance out of the total expression variance, and truly variable genes will usually show a lower correlation with the cell depth compared to housekeeping genes with similar expression average. We therefore compute an empirical trend *r*(*u*_*g*_) using the median 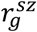 correlation in a moving window of 100 genes ordered by total gene expression *u*_*g*_. We then define the normalized depth scaling as 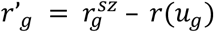. Finally, we select genes with sufficiently high *u*_*g*_ and *r*′_*g*_ < *T*_*gr*_ (typically, *T*_*gr*_ = −0. 1). We note this approach for selecting feature genes may be biased against genes that are enriched for cell types with high *u*_*i*_ distribution, and that careful manual analysis of the selected feature genes is recommended in all cases.

